# Patterns of genetic differentiation imply distinct evolutionary histories of the sibling mosquito species *Anopheles messeae* and *Anopheles daciae* in Eurasia

**DOI:** 10.1101/2022.11.23.517664

**Authors:** Ilya I. Brusentsov, Mikhail I. Gordeev, Andrey A. Yurchenko, Dimitriy A. Karagodin, Anton V. Moskaev, James M. Hodge, Vladimir A. Burlak, Gleb N. Artemov, Anuarbek K. Sibataev, Norbert Becker, Igor V. Sharakhov, Elina M. Baricheva, Maria V. Sharakhova

**Affiliations:** Department of Entomology, Virginia Polytechnic Institute and State University and Fralin Life Sciences Institute, Blacksburg, VA, USA; Laboratory of Cell Differentiation Mechanisms, Institute of Cytology and Genetics, Novosibirsk, Russia; Department of General Biology and Ecology, Moscow Region State University, Moscow, Russia; Laboratory of Ecology, Genetics, and Environmental Protection, Tomsk State University, Tomsk, Russia; Department of General Biology and Genomics, L.N. Gumilyov Eurasian National University, Astana, Kazakhstan; Department of Agricultural Biology, Tomsk State University, Tomsk, Russia; Center for Organismal Studies, University of Heidelberg, Heidelberg, Germany; German Mosquito Control Association, Speyer, Germany

**Keywords:** *Anopheles*, mosquito, inversion polymorphism, phylogeography, evolutionary history, population genetics

## Abstract

Detailed knowledge of phylogeography is important for control of mosquito species involved in transmission of human infectious diseases. *Anopheles messeae* is a geographically widespread and genetically diverse dominant vector of malaria in Eurasia. A closely related sibling species, *An. daciae*, was distinguished from *An. messeae* based on a few nucleotide differences in its ribosomal DNA. However, the mechanisms of speciation and their evolutionary histories are poorly understood. Here, we performed a large-scale population genetics analysis of 3694 mosquitos from Eurasia to understand the species divergence, diversity, and population structure using the Internal Transcribed Spacer 2 of ribosomal DNA for species identification and frequencies of 11 polymorphic chromosomal inversions as genetic markers. The study revealed striking differences in the geographical distribution of the sibling species. The largest genetic differences between *An. messeae* and *An. daciae* were detected in the X sex chromosome suggesting that this chromosome plays a role in speciation. The frequencies of autosomal inversions differed significantly between the species, strongly supporting a restricted gene flow. The clinal variability of some inversion frequencies was revealed in both species implicating their possible involvement in climate adaptations. Statistical analysis of inversion polymorphism clearly distinguished two clusters associated with the two species and demonstrated much higher genetic diversity within *An. messeae*. Overall, the frequencies of hybrids in all locations were extremely low with the exception of several southeastern populations, where putative hybrids were abundant. Thus, the pattern of genetic differentiation implies dramatic differences in geographic distribution, population structure, and evolutionary histories of the sibling species *An. messeae* and *An. daciae*.

## Introduction

Phylogeography is a rapidly developing discipline that analyzes the genealogy of lineages within the context of their geographical distribution (Emerson & Hewitt, 2005). It is employed to better understand the evolutionary histories of populations, subspecies, and species. Application of different molecular markers and large numbers of specimens from natural populations, together with modern analytical tools, provides important insights into the distribution of genetic diversity around the globe. Chromosomal inversions play an important role in the evolution of living organisms (F. J. Ayala & Coluzzi, 2005) and, for a long time, they served as unique markers of population genetics and phylogenetics in species with well-developed polytene chromosomes, such as *Drosophila* (Krimbas & Powell, 1992) and malaria mosquitoes from the *Anopheles* genus (Coluzzi, Sabatini, Della Torre, Di Deco, & Petrarca, 2002; Stegniy, 1991). Here, we argue that using molecular markers and polymorphic chromosomal inversions in combination with modern statistical methods helps us to better understand the evolutionary histories of well-known and newly described species of malaria mosquitoes.

Among different infectious diseases, malaria is traditionally considered the most dangerous disease transmitted to humans by mosquitoes (Rougeron et al., 2022). Despite extensive efforts to eradicate malaria, global climate change, human migration, political instability, and the presence of competent malaria vectors have increased the risk of malaria importation and transmission in regions where it was previously eliminated (Chretien et al., 2015; Kulkarni, Duguay, & Ost, 2022; Rossati et al., 2016; Sainz-Elipe et al., 2010). Thus, understanding the phylogeography of malaria mosquitoes is important for the prediction of malaria propagation and for developing effective and adequate strategies for eradication of this disease. Moreover, mosquitoes represent an excellent model to study genetic mechanisms of species evolution because mosquito biology, ecology, and molecular genetics are well-developed disciplines (Powell, 2018).

The most dangerous vectors of malaria in the Northern Hemisphere belong to the Maculipennis group (Sinka et al., 2010). The major vector of malaria in Europe, *Anopheles maculipennis*, Meigen, 1818, was originally considered as a single species until the phenomenon of, so called, “anophelism without malaria” was observed (Hackett, 1937; Hackett & Missiroli, 1935). It was shown that in some parts of Europe, malaria was absent despite the presence of malaria vectors. This phenomenon led to the conclusion that *An. maculipennis* represents a complex of species with different abilities to transmit malaria. The presence of multiple species within the complex was supported by hybridization experiments (Kitzmiller, Frizzi, & Baker, 1967; Stegniy, 1980; Stegniy, Sichinava, & Sipovich, 1984) and by morphological differences of their egg chorion patterns (Gutsevich, Monchadsky, & S, 1970). Later, chromosomal analysis became an additional robust tool for species identification, distinguishing the taxonomic status of a species, and for population analysis. For example, fixed chromosomal differences discriminated *An. maculipennis* and *An. messeae* Falleroni, 1926 (Stegniy, Pestryakova, & Kabanova, 1973) and *An. sacharovi* Favre, 1903 and *An. martinius* Shingarev, 1926 (Stegniy, 1976). A new species from the Maculipennis group, *An. beklemishevi*, Stegniy and Kabanova 1976, was specifically identified based on fixed chromosome differences (Stegniy & Kabanova, 1978; Stegniy, Kabanova, & Novikov, 1976). Using the Internal Transcribed Spacer 2 (ITS2) of ribosomal DNA (rDNA) as an additional molecular marker for species diagnostics resulted in the identification of several new species: *An. artemievi* Gordeyev, Zvantsov, Goryacheva, Shaikevich & Yezhov, 2005, *An. daciae* Linton, Nicolescu & Harbach, 2004, and *An. persiensis* Linton, Sedaghat & Harbach, 2003. Currently, the Maculipennis group comprises eleven Palearctic species, four of which, *An. atroparvus*, Van Thiel, 1972, *An. labranchiae*, Falleroni 1926, *An. sacharovi*, and *An. messeae* are considered dominant vectors of malaria in Eurasia (Sinka et al., 2010).

*An. messeae* is the most geographically widespread and genetically diverse species among the other dominant malaria vectors from the Maculipennis group. Distribution of the species extends from Ireland in the west to the Amur River region in the east and from Scandinavia and Yakutia in the north to Iran and northern China in the south (Gornostaeva & Danilov, 2002; Sinka et al., 2010; Zvantsov, Gordeev, Goriacheva, & Ezhov, 2014). Despite being largely zoophilic, *An. messeae* actively feeds on humans in cases of animal deficiency (Fyodorova et al., 2006). *An. messeae* is the primary malaria vector of *Plasmodium vivax* malaria in Russia (Beklemishev, 1948; Daskova & Rasnicyn, 1982). As the most genetically diverse species among the Maculipennis subgroup, *An. messeae* carries five widely spread chromosomal inversions X1, X2, 2R1, 3R1, and 3L1, located on four chromosomal arms (Kabanova, Kartashova, & Stegniy, 1972). The details of inversion polymorphism in natural populations of *An. messeae* were intensively investigated by a group of researchers led by V. Stegniy from 1970-1990s (Stegniy, 1991). These studies demonstrated a complex structure of *An. messeae* populations along the range of the species distribution. It was shown that inversions played an important role in climatic adaptation of the species in latitudinal and meridional directions (Stegniy, Kabanova, Novikov, & Pleshkova, 1976). For example, a latitude cline was described for the 2R1 inversion where the inverted variant was more abundant in northern populations, suggesting that this inversion could be involved in adaptation to cold temperatures and to successful overwintering. In contrast, frequencies of the X1 and 3R1 inversions displayed a west-east longitude cline with higher frequencies of the inverted variants found in eastern populations. Involvement of the chromosomal inversion in local adaptations of the species (Stegniy, Novikov, Pleshkova, & Kabanova, 1978) and seasonal (Kabanova, Stegniy, & Luzhkova, 1973) and temporal dynamics (Pleshkova, Stegniy, Novikov, & Kabanova, 1978) in several geographical locations was also observed. Overall, these studies led to conclusions about the stable state of the inversion polymorphisms in natural populations of *An. messeae* (Stegniy, 1983a) and the dependence of chromosomal variability on landscape-climatic zones (Gordeev & Moskaev, 2016). Thus, the idea of a potential role of chromosomal inversions in genetic divergence among *An. messeae* populations was rejected by these studies.

The hypothesis that combinations of different chromosomal inversions in natural populations of *An. messeae* represent two distinct chromosomal complexes was first introduced in 1979 by Y. Novikov and V. Kabanova (Novikov & Kabanova, 1979). They showed that frequencies of some inversion combinations were higher or lower than expected from random mating, suggesting the presence of linkage disequilibrium between them. For example, the standard karyotype X0 was more frequently associated with standard karyotype variants 2R0, 3R0, and 3L0, whereas the inverted karyotypes X1 and X2 were associated with 2R1, 3R1, and 3L1 inversions (Gordeev & Stegniy, 1987; Novikov & Kabanova, 1979; Stegniy, 1983b). Later studies demonstrated that these chromosomal complexes varied by female fecundity (Gordeev & Stegniy, 1987), larval diet (Gordeev & Troshkov, 1990), sensitivity to the toxins of *Bacillus thuringiensis* (Gordeev & Burlak, 1991), rate of survivorship of adults and larvae under laboratory conditions (Gordeev & Perevozkin, 1995), and rate of larval survivorship under selection pressure related to predators (Gordeev, Perevozkin, & Luk’yantsev, 1997; Gordeev & Sibataev, 1995) and parasites (Burlak & Gordeev, 1998). In 2001, these chromosomal complexes were referred to as cryptic genetically isolated forms or as species named “A” and “B” (Novikov & Shevchenko, 2001).

As mentioned above, in 2004, independent of these studies, a new Maculipennis species, *An. daciae*, was differentiated from *An. messeae* based on five ITS2 nucleotide substitutions and morphological differences at the egg stage (Nicolescu, Linton, Vladimirescu, Howard, & Harbach, 2004). Later, sequencing of ITS2 indicated that chromosomal form “A” of *An. messeae* is synonymous with *An. daciae* (Vaulin & Novikov, 2012). Additional geographical studies discovered *An. daciae* in Germany (Kampen, Schäfer, Zielke, & Walther, 2016; Kronefeld, Werner, & Kampen, 2014; Weitzel, Gauch, & Becker, 2012), the United Kingdom (Danabalan, Monaghan, Ponsonby, & Linton, 2014), Poland (Rydzanicz, Czułowska, Manz, & Jawień, 2017), the Czech Republic, Slovakia (Blažejová et al., 2018), Serbia (Kavran et al., 2018), Finland (Culverwell, Vapalahti, & Harbach, 2020), Sweden (Lilja, Eklöf, Jaenson, Lindström, & Terenius, 2020), Belgium (Smitz et al., 2021), and Italy (Calzolari et al., 2021). Significant differences in the frequencies of chromosomal inversions between *An. messeae* and *An. daciae* were determined from three populations from the Moscow region (Naumenko et al., 2020). This study also indicated that the inverted variant X1 was fixed in *An. messeae* populations while the standard X0 and inverted X1 karyotypes were segregated in *An. daciae* populations from the same locations. Whole genome sequencing analyses demonstrated genome-wide differentiation between the species, which was especially pronounced on chromosome X. However, some authors doubt the separate species status of *An. messeae* and *An. daciae* and consider the genetic differences between them as intraspecies polymorphism within *An. messeae* (Bertola, Mazzucato, Pombi, & Montarsi, 2022; Bezzhonova & Goryacheva, 2008; Calzolari et al., 2021; Stegniy, Pishchelko, Sibataev, & Abylkassymova, 2016).

The goal of this study is two-fold: 1) to evaluate the potential role of chromosomal inversion in the evolution of the malaria vector-species *An. messeae* and *An. daciae*, and 2) to better understand their evolutionary histories in Eurasia. For the first time, chromosomal inversions were analyzed in *An. messeae* and *An. daciae* separately in multiple populations across Eurasia. Using polymorphic inversions as genetic markers, the detailed statistical analyses of divergence, diversity, and population structure of the two species was performed. In this paper, we use the taxonomic nomenclature proposed by Nicolescu and Linton (Nicolescu et al., 2004) and consider *An. messeae* and *An. daciae* as separate sibling species because this assumption agrees with our own findings.

## Materials and Methods

### Sample collection and material preservation

A total of 3694 mosquito larvae were collected from 2017-2019 in 28 locations across 20 regions in Germany, Kazakhstan, Latvia, and Russia (Fig. 1, Table S1). The collections were carried out in late July – early August, when both species are abundant in the natural populations (Czajka, Weitzel, Kaiser, Pfitzner, & Becker, 2020). *Anopheles* larvae were collected using the dipping method. Each individual mosquito larva was numbered and divided into two parts: head with thorax and abdomen. These dissected parts were placed into separate tubes for further analysis. Heads with thoraxes were kept in Carnoy’s solution (1:3 acetic acid: ethanol) and abdomens were placed in 70% ethanol. Additionally, we included data from three locations in the Moscow region (Noginsk, Novokosino, and Yegoryevsk), which had been published earlier (Naumenko et al., 2020).

**Fig. 1.**
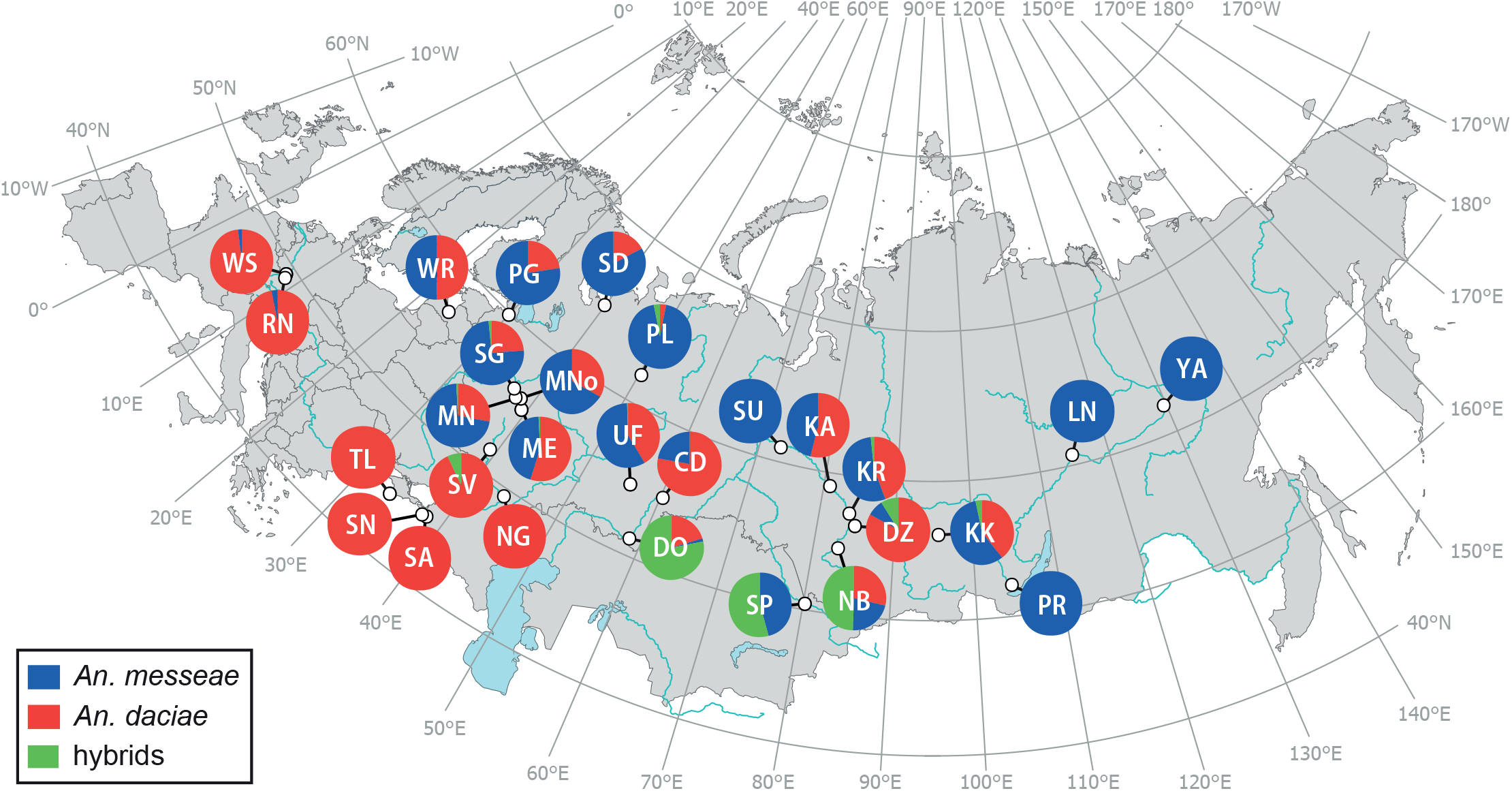
Compositions of *Anopheles messeae, Anopheles daciae*, and their hybrids in different locations in Eurasia. The proportions of *An. messeae, An. daciae*, and their hybrids are shown as pie charts with different colors for each site. Collection sites are indicated as: CD – Desyatiletie, DO – Dombarovka, DZ – Dzerzhinskoe, KA – Kargasok, KK – Kansk, KR – Krivosheino, LN – Lensk, ME – Yegoryevsk, MN – Noginsk, MNo – Novokosino, NB – Berdsk, NG – Novogrigor-yevskaya, PG – Petergof, PL – Palevitsy, PR – Irkutsk, RN – Reinhausen, SA – Abinsk, SD – Severodvinsk, SG – Solnechnogorsk, SP – Semey, SN – Novorossiysk, SU – Surgut, SV – Voronezh, TL – Tylovoe, UF – Ufa, WR – Riga, WS – Speyer, YA – Yakutsk. Overall, *An. messeae* is considered as a north-eastern species and *An. daciae* as south-western species. Intriguingly, a high number of hybrids were found in some southern locations of the explored area.

### Genotyping

For genotyping, the abdomen of each sample was homogenized in liquid nitrogen and genomic DNA was extracted using the standard protocol for the Qiagen DNeasy Blood and Tissue Kit (Qiagen, Germantown, MD, USA). DNA elution was performed in 100 μl of water. ITS2 from rDNA was amplified using the forward universal primer its2_mdir 5’-GCTCGTGGATC-GATGAAGAC-3’ (with modifications from Djadid et al., 2007), Tm=57°C or its2_vdir 5’-TGTGAACTG-CAGGACACATG-3’ and the reverse primer its2_rev 5’-ATGCTTAAATTTAGGGGGTAGTC-3’, Tm=54°C with modifications (Proft, Maier, & Kampen, 1999). PCR with a combination of the its2_mdir and its2_ rev primers produced amplicons of 477 bp (435 bp for its2_vdir and its2_rev) in length for *An. messeae* and *An. daciae*, 464 bp for *An. maculipennis* (422 bp forits2_vdir and its2_rev), 479 (437 bp for its2_vdir and its2_rev) for *An. atroparvus* and >800 bp for *An. beklemishevi*. The HotStarTaq Plus Master Mix Kit (Qiagen, USA) was used for PCR. The PCR mixture contained a total volume of 20 μl of ~40 ng of DNA, 0.5 μM of each forward and reverse primer, and 10 μl of 2× HotStarTaq Plus reaction-mix. PCR was performed using a thermal cycler (Eppendorf, Hauppauge, NY, USA) with the following conditions: initial denaturation at 95°C for 5 min, followed by 25 cycles of 95°C for 15 sec, 58°C for 30 sec, and 72°C for 30 sec, and a final extension step at 72°C for 5 min. The reaction mix was then placed on hold at 4°C. Amplicons were visualized using gel electrophoresis in 2% agarose gel.

For DNA sequencing, amplicons were purified with a Wizard^TM^ PCR Clean Up kit (Promega, Fitchburg, WI, USA). PCR products were sequenced using the BigDye Terminator v3.1 Cycle Sequencing Kit (Thermo Fisher Scientific, USA) from suitable forward or reverse primers, and sequencing reactions were purified by chromatography on Sephadex G-50 (Sigma, USA) and analyzed in the SB RAS Genomics Core Facility (http://sequest.niboch.nsc.ru/). All specimens of *An. daciae, An. messeae*, and *An. atroparvus* were genotyped by Sanger sequencing. Species of *An. maculipennis* and *An. beklemishevi* were mostly identified by karyotyping but some were verified by sequencing. A total of more than 2600 specimens were successfully genotyped and analyzed using SeqScape v 3.0 software (Thermo Fisher Scientific, Haverhill, MA, USA). The ITS2 sequence of *An. messeae* AY648982 (Nicolescu et al., 2004) was used as a reference. Basecalling was performed using KB basecaller with mixed base identification enabled. Nucleotide positions in ITS2 of *An. messeae* s.l. were given in correspondence to AY648982 (Nicolescu et al., 2004). The previously described SNPs, in positions 150, 211, 215, 217, 412, and 432, distinguished the ITS2 sequences of *An. messeae* from *An. daciae* (Naumenko et al., 2020) and were investigated in detail. For this, data from basecalled.ab1 files were extracted by Perl script based on Bio::Trace::ABIF v 1.06 perl module (http://search.cpan.org/~vita/Bio-Trace-ABIF-1.06/). Peaks were visualized and analyzed in Mathematica v 12 software (Wolfram Research, Champaign, IL, USA) for base assignment based on the Sign test of peak intensity and 3 background signal intensities from each side of the peak. To verify the appropriateness of the nucleotide base assignment in these positions, diagnostic SNPs were manually analyzed. Only species-specific substitutions in positions 412 and 432, as G, G for *An. messeae* and A, C for *An. daciae* were used for diagnostics of the species and their hybrids. Samples with unusual 412 and 432 base combinations were considered unconventional and were excluded from the analysis.

### Karyotyping

Salivary glands were dissected from the larval thorax for preparation of polytene chromosomes and chromosome preparations were made by the standard lac- to-aceto-orcein method (Kabanova et al., 1972). Polytene chromosomes were visualized using an Eclipse E200 light microscope (Nikon, BioVitrum, Moscow, Russia). Specimens were karyotyped using chromosome maps for the salivary glands of *An. messeae* and *An. daciae* (Artemov et al., 2021; Stegniy, Kabanova, & Novikov, 1976). A total of 11 chromosomal inversions — X1, X2, X3, X4, 2R1, 2R3, 2R4, 3R1, 3R2, 3L1, and 3L4 – were identified and considered in this study (Table 1). The karyotypes of each specimen were described for the whole chromosomal complement. A total of 3133 samples were successfully karyotyped.

**Table 1.**
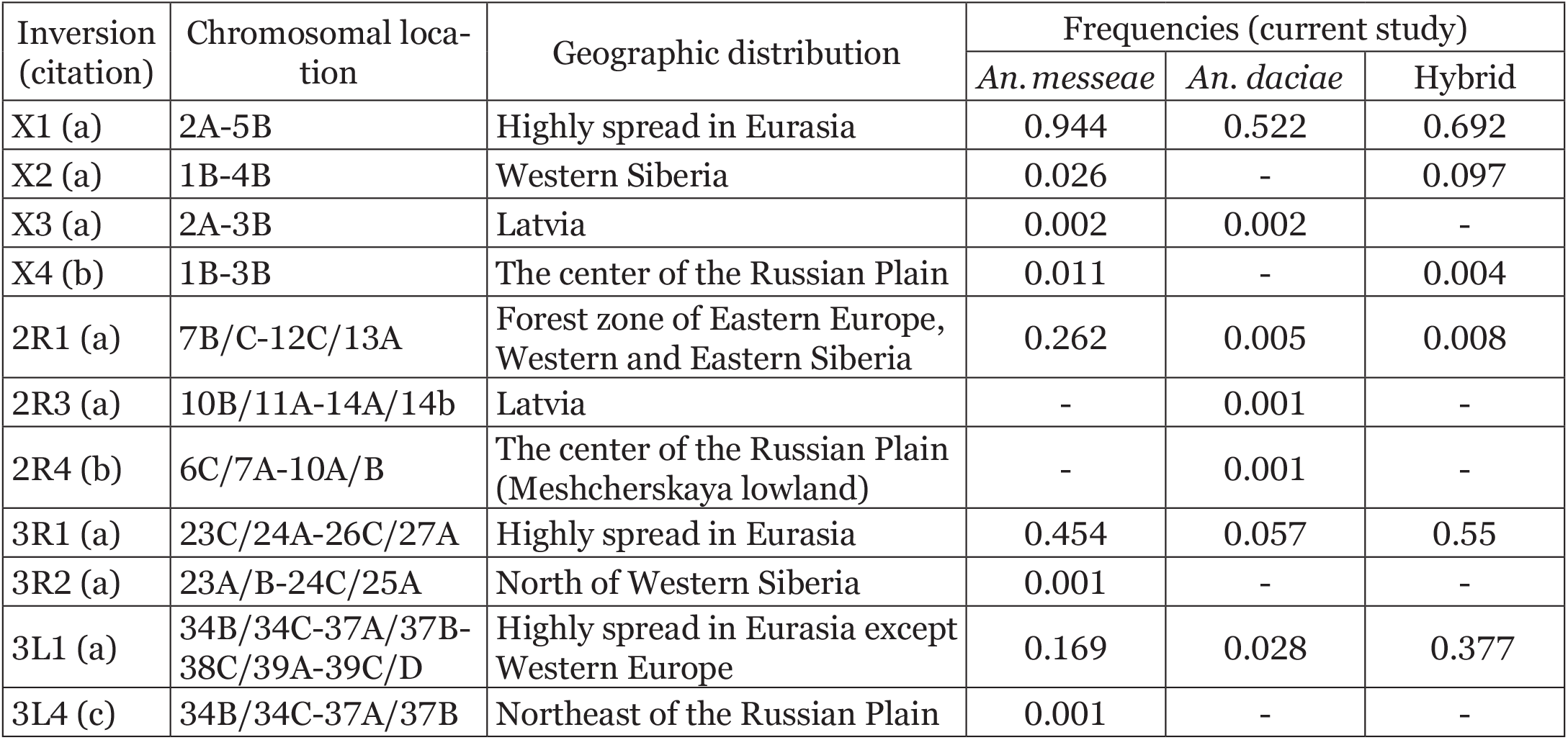
Chromosomal inversions in *Anopheles messeae* and *Anopheles daciae*. Inversions were first described in: a) Stegniy, Kabanova, & Novikov, 1976; b) Naumenko et al., 2020; and c) this study.

### Statistical analysis of chromosomal inversion frequencies

Samples of *An. messeae* and *An. daciae* that were genotyped and karyotyped (Table S2) were used for statistical analysis. Chromosomal inversions were utilized as genetic markers for statistical analysis using R packages (R_Core_Team, 2021). Species and population differentiations were analyzed using the «poppr» package in R (Kamvar, Tabima, & Grünwald, 2014). Genotypic diversity was assessed by three parameters. First, the richness of multilocus genotypes (MLG) was assessed by the Menkhinik index (Menhinick, 1964), demonstrating a combination of all chromosomal variants for each individual mosquito. Second, the Simpson Index was calculated to estimate the probability of two random mosquitoes having a different genotype (Simpson, 1949). For elimination of the error related to differences in sample size, an adjustment was made by multiplying them by n/(n-1), where n is the count of mosquitoes in the sample. Third, the Shannon-Wiener diversity index (Shannon, 1948) quantified the uncertainties associated with predicting the MLG in the next individual based on already identified MLGs.

Deviations from the Hardy-Weinberg equilibrium were assessed using the exact test of Hardy-Weinberg equilibrium (Wigginton, Cutler, & Abecasis, 2005) implemented in the Hardy-Weinberg R package (Graffelman, 2015) for each autosome for all populations. Genetic differentiation (Hedrick’s G’st) between species by chromosomes was calculated using the “mmod” R package (Winter, 2012).

Assessment of the genetic diversity in the studied species and population differentiation was performed using Analysis of Molecular Variance (AMOVA) (Excoffier, Smouse, & Quattro, 1992; Michalakis & Excoffier, 1996). Countries or federal districts and sampling sites were used to partition the data into different stratifications. The pairwise Fst values (Weir & Hill, 2002) between the studied populations were calculated using the “BEDASSLE” package (Gideon, 2014). To assess isolation by distance, a correlation test between pairwise Fst and the pairwise distance between populations was carried out. The distances were calculated using the formula: RE · arccos(sin(x_1_) · sin(x_2_) + cos(y_1_) · cos(y_2_) · cos(|y_1_-y_2_|)), where (x, y) are the location coordinates, and RE – radius of Earth (6373 km.). A correlation test between Fst and distance was carried out to assess the isolation by distance.

Hierarchical clustering analysis and Principal Component Analysis (PCA) were performed using the frequencies of autosomal chromosomal inversions on the population level to infer and visualize the relationship between the species and populations (Package for R “stats” (R_Core_Team, 2021)).

## Results

### Geographical distribution of the Maculipennis species in Eurasia

In this study, we analyzed the composition of Maculipennis species in 28 locations in 20 regions in four countries in Eurasia (Fig. 1, Table S1) using ITS2 analyses and karyotyping. The species were differentiated based on ITS2 length or sequence. For example, *An. beklemishevi* was recognized based on the length of the PCR product of ITS2, which is significantly longer than in other species. Because the ITS2 length in *An. daciae, An. atroparvus*, and *An. maculipennis* differ from that of *An. messeae* by 0, 2, and 13 bp, respectively, these species were recognized by sequence analyses (Kampen, 2005; Naumenko et al., 2020). As mentioned before, *An. daciae* was originally described based on five chromosomal substitutions in positions 211, 215, 217, 412, and 432 (Nicolescu et al., 2004). However, a later study found heterogeneity in the first three nucleotide positions 211, 215, 217 – W(A+T), W(A+T), and Y(C+T), respectively – in each specimens of *An. daciae* from the Moscow region and an additional heterogenic nucleotide, M(A+C), in position 150 of *An. messeae* (Naumenko et al., 2020). Thus, reliable species diagnostics can be conducted only based on species-specific nucleotides in positions 412 and 432. Our current study confirmed previous observations regarding heterogeneity of nucleotides 211, 215, 217 in *An. daciae* and nucleotide 150 in *An. messeae*.

The representation of diagnostic nucleotide variants in ITS2 sequences in all samples is shown in Fig. 2. For *An. messeae*, the main variant MTTCGG (95%) was observed earlier in the populations in the Moscow region (Naumenko et al., 2020). The second most common variant (3.7%) was without the heterogeneity at the 150 nucleotide (CTTCGG). Moreover, most of the samples, 18 out of 25, with this variant were found in the two eastern populations of Yakutsk and Lensk, where *An. messeae* was present without an admixture with other species. Two additional combinations containing a heterogeneous nucleotide K (T+G) at position 211 were found in 3 populations: Severodvinsk, Yakutsk, and Krivosheino, at only 1.5%. Significantly greater diversity of 10 variants was determined in *An. daciae* samples in diagnostic substitutions. They were associated with heterogeneous nucleotides in positions 211, 215, and 217, which contained various ratios of the major and additional nucleotides over a wide range. The major variant was CWWYAC (81%), containing heterogeneous substitutions at all three positions, but variants containing one, two, and no heterogeneous substitutions were also identified (Fig. 2). The most diverse were combinations of diagnostic nucleotides that were found in hybrids of *An. messeae* and *An. daciae*. In addition to all these variants, we discovered unconventional sequencing results in *An. messeae* and *An. daciae* having species-specific substitutions in the diagnostic positions 412 and 432 diagnostic positions in 49 (2.53%) mosquito samples. Individuals with such nucleotide composition were excluded from further analysis.

**Fig. 2.**
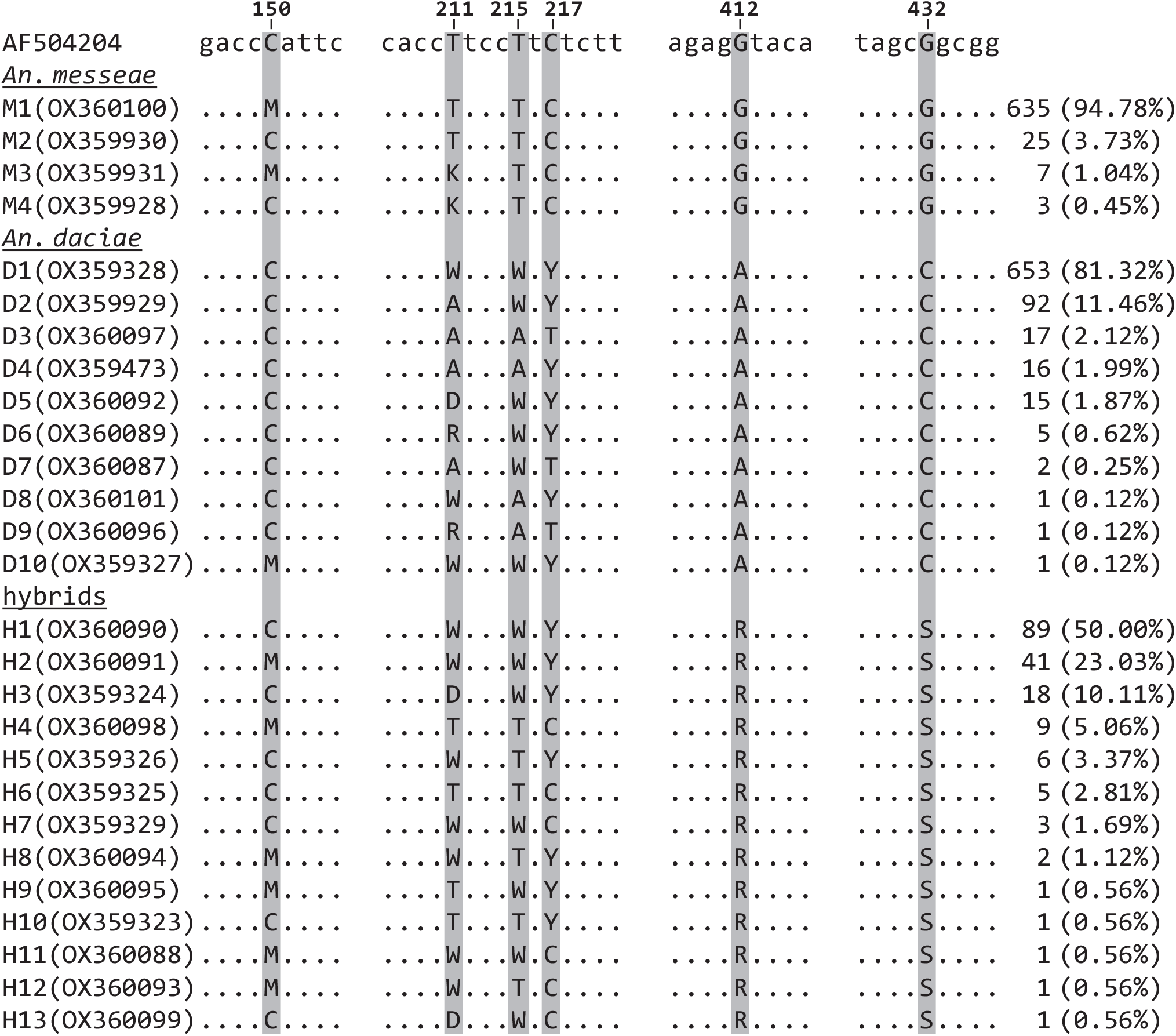
Variations in five diagnostic nucleotides of ITS2 sequences in *Anopheles messeae, Anopheles daciae*, and their hybrids. The coordinates of the diagnostic nucleotides in the ITS2 sequence of *An. messeae* (AY648982) are shown above according to the primary description of the species (Nicolescu et al., 2004). The designation of variants of diagnostic nucleotides and their accession numbers obtained in this study are indicated on the left. Nucleotides corresponding to the reference ones are shown as dots, but other than the reference ones are shown as letters. The amount and frequencies of each variant inside a species are shown on the right. The number of variations was higher in *An. daciae* and hybrids than in *An. messeae*.

Our study revealed the presence of five species among the 11 species of the Palearctic members of the Maculipennis group in our study locations: *An. atroparvus, An. beklemishevi, An. daciae, An. maculi-pennis*, and *An. messeae*. *An. messeae* and *An. daciae* were the most abundant species in the studied regions. The geographical distribution of these two species widely overlapped in the central part of Eurasia: in Latvia and in the central part of European Russia, Ural, and Western Siberia (Fig. 1, Table S3). However, in the peripheral locations, the species composition varied from the complete absence of *An. messeae* in the southern regions of Russia (Abinsk, Novorossiysk, Voronezh, Novogrigoryevskaya, Dombarovka, and Tylovoe) and Germany (Reinhausen, Speyer), to the complete absence of *An. daciae* in the northern regions of Russia (Palevitsy and Surgut), Eastern Siberia (Irkutsk, Lensk, Yakutsk), and Kazakhstan (Semey) (Fig. 1, Table S3). Among all 1932 individual mosquitoes analyzed based on ITS2 sequencing, 179 individuals, or 9.27%, were considered to be hybrids between *An. daciae* and *An. messeae*. The ITS2 sequences in these specimens demonstrated double peaks in the last two species-diagnostic positions. Interestingly, in most locations where distributions of the species were overlapped, hybrids represented only ~1% of the specimens, but a significant number of hybrids between *An. daciae* and *An. messeae* were found in four locations: 53.54% in Semey, Kazkhstan 78.13% in Dombarovka, Orenburg region, and in two locations in the south of Western Siberia with 49.43% in Berdsk and 8.89% in Dzerzhinskoe (Fig. 1, Table S3). Moreover, in addition to these hybrids, only *An. messeae* or *An. daciae* were found in Semey and in Dombarovka, respectively.

As mentioned above, three other species, *An. maculipennis*, *An. atroparvus*, and *An. beklemishevi*, were found in several locations in Russia. *An. maculipennis* was widespread in two southern locations close to the Black Sea with 67% in Tylovoe and 36% in Novorossiysk. A total of 20% of this species was found in another European location in the Moscow region (Yegoryevsk). Another species, *An. beklemishevi* was present in the Tomsk region of Western Siberia at the level of 88% in Kargasok and 15% in Krivosheino, in Eastern Siberia at the level of 10% in Lensk, and in the Northern European part of Russia at the level of 5% in Palevitsy. Only two specimens of *An. atroparvus* were found in Tylovoe, Crimea.

Overall, our study clearly demonstrated differences in geographical distribution between *An. messeae* and *An. daciae*, identifying them as northeastern and southwestern species, respectively. In addition, in our study, *An. beklemishevi* was identified as a northeastern species and *An. maculipennis* as a southwestern species.

### Distinct patterns of inversion distribution in An. messeae and An. daciae

A polytene chromosomal compliment in all *Anopheles* species, including *An. messeae* and *An. daciae*, is composed of 3 chromosomes (X, 2, and 3) comprising 5 chromosomal arms: X, 2R, 2L, 3R, and 3L (Fig. 3). Because one of the arms in the sex chromosome is absent in a polytene chromosome compliment, we indicated it as X, but not as XL or XR (Artemov et al., 2021). The Y chromosome is heterochromatic and is absent in the polytene chromosome compliment. This study included 670, 803, and 178 individuals of *An. messeae*, *An. daciae*, and their hybrids, respectively. We found the common inversions X1, 2R1, 3R1, and 3L1, inversions X2, X3, 2R3, 3R2, which have been previously described as endemic (Stegniy, Kabanova, & Novikov, 1976), rare inversions X4 and 2R4 (Naumenko et al., 2020), and a newly described rare inversion 3L4 in 28 locations across Eurasia (Table 1). The positions of the breakpoints for these

**Fig. 3.**
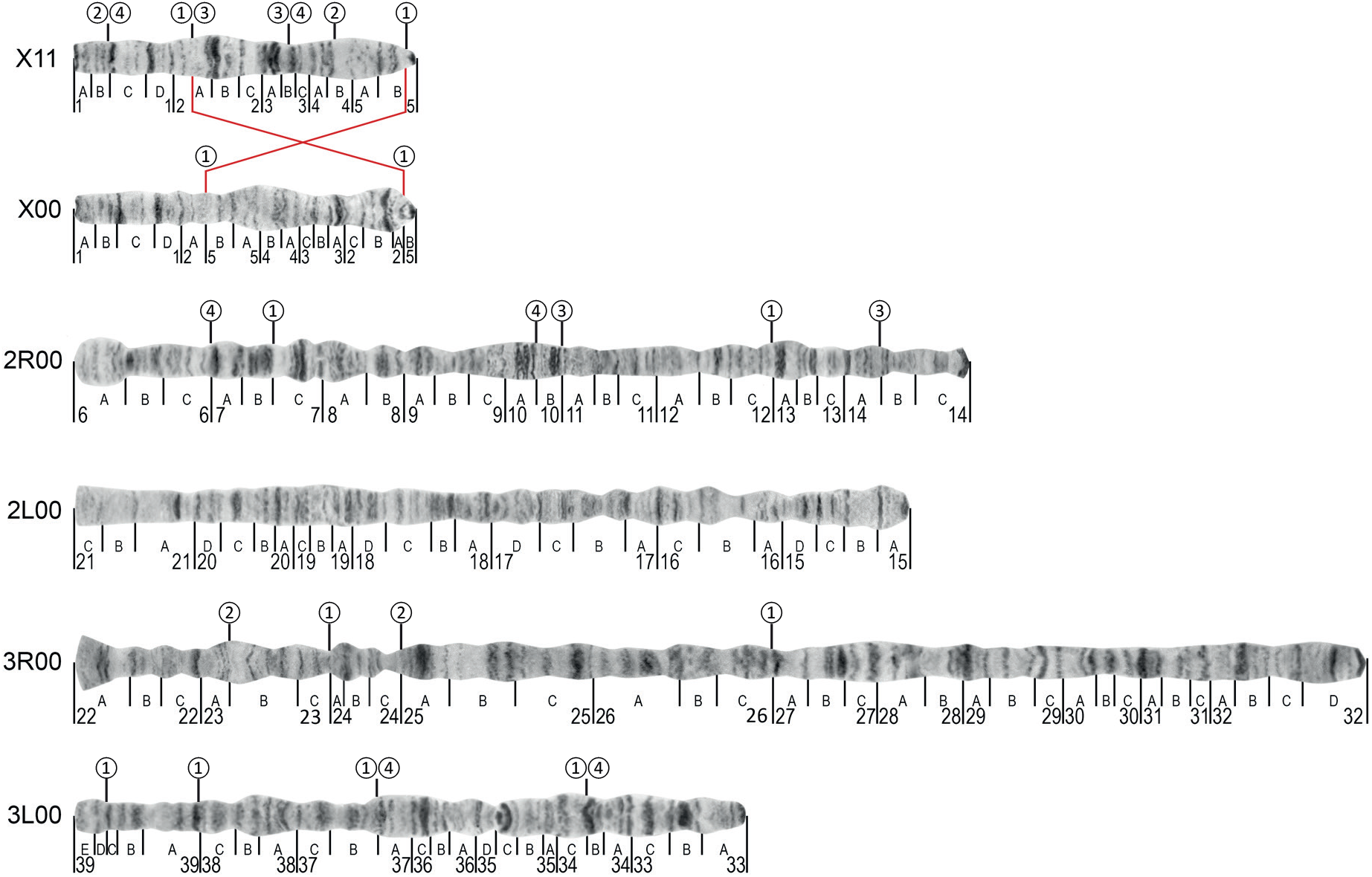
A cytogenetic map of chromosomal inversions in *Anopheles messeae* and *Anopheles daciae*. Standard X00 and inverted X11 karyotypes are shown for the sex chromosome. Standard chromosome arrangements are shown for autosomal arms 2R00, 2L00, 3R00, and 3L00. Inversion breakpoints are indicated by numbers inside of the circles above the chromosomes. Inversion X1 is additionally shown by red brackets. Numbered divisions and lettered subdivisions of the chromosomes are shown below the chromosomes. The order of the divisions on chromosome 1 is based on inverted arrangement X11 according to the previous published chromosomal map (Artemov et al. 2021). A total of 11 chromosomal inversions are shown. All chromosomal inversions on chromosome X are based on the inverted arrangement X11.

inversions are shown in Fig. 3. According to the common inversion nomenclature for *An. messeae*, we referred to standard or “not inverted” homokaryotypes as a 00 variant and inverted homokaryotypes as a 11 or 22 variant; heterokaryotypes were referred to as 01, 02, 03, or 04 variants. Inversions on the sex chromosome X in hemizygous males were referred as 0, 1, 2, 3, and 4 variants. Homokaryotypes 22, 33, and 44 were absent in this study.

Our analyses demonstrated that some chromosomal inversions were species-specific and frequencies of other chromosomal inversions varied between *An. messeae, An. daciae*, and their hybrids (Fig. 4, Table 1). A standard karyotype X0 was found in *An. messeae* at an extremely low frequency of 0.02, in most cases in hemizygote males or heterozygote females, whereas inverted variant X1 was the most prevalent in this species. In contrast, in *An. daciae*, the frequency of the X0 variant was 0.48. In hybrids, the frequency of this variant was 0.21. Inversion X2 was detected only in *An. messeae* populations in Western Siberia and Eastern Kazakhstan but was absent in European regions. The frequency of this inversion was 0.03. Inversion X4 was rare with a frequency of 0.01 and it was found only in *An. messeae* in Moscow populations. This chromosomal variant was observed only in hemizygote males and heterozygote females. In hybrids, inversions X2 and X4 were also found in Kazakhstan and in Moscow, respectively. The rare inversion X3 was only found in Latvia in both *An. messeae* and *An. daciae* with a frequency less than 0.01 in both species (Fig. S1). Moreover, in all cases, this inversion was present in hemizygote males in *An. messeae* but in heterozygote females in *An. daciae*. Autosomal inversion 2R1 was highly abundant in *An. messeae* (0.26) but found at extremely low frequency (0.01) in *An. daciae* and hybrids where it was found mostly as heterozygotes. The inverted autosomal variants 3R1 and 3L1 were present in both *An. messeae* and *An. daciae* but their frequencies were significantly higher in *An. messeae* and hybrids (Table 1). In contrast, the rare chromosomal inversions 2R4 and 2R3 were found as heterozygotes only in *An. daciae*.

**Fig. 4.**
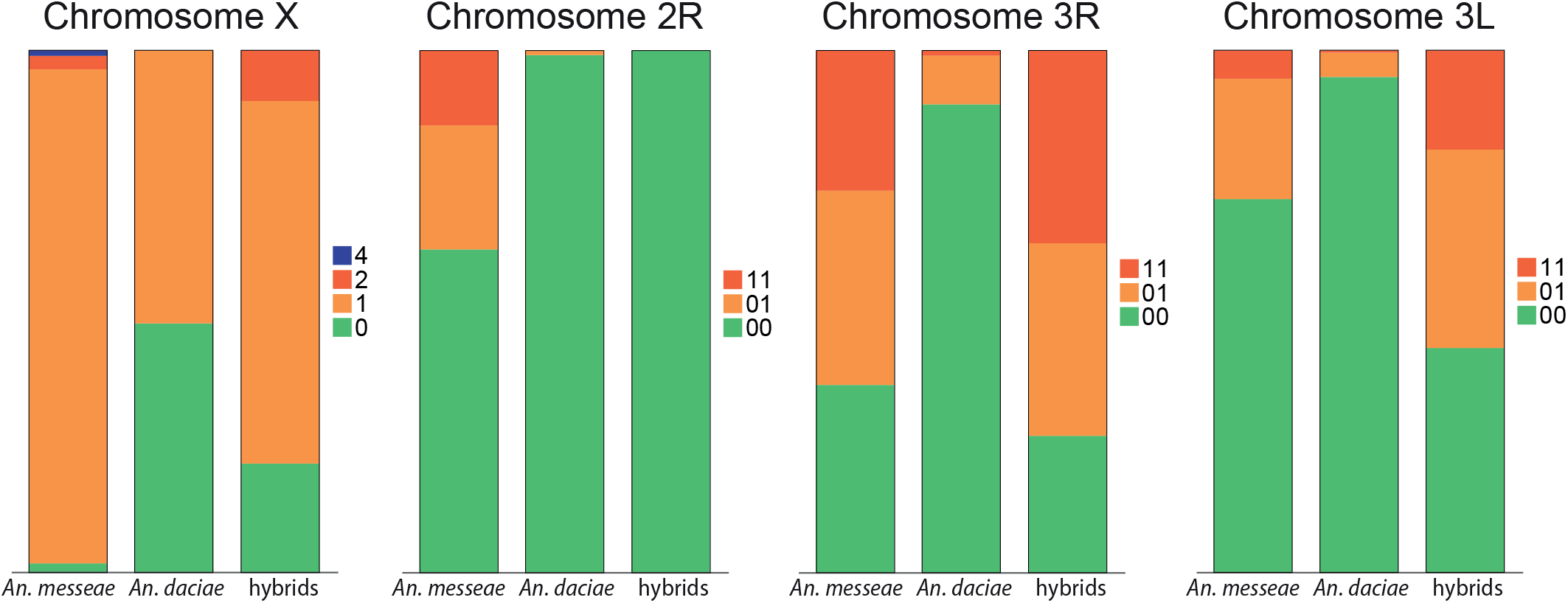
The frequencies of the chromosomal inversion variants in *Anopheles messeae, Anopheles daciae*, and their hybrids. Proportions of standard, inverted, and heterozygote arrangements are shown by different colors. Frequencies of allelic chromosome variants for chromosome X and frequencies of diploid genotypes for autosomes are shown. *An. messeae, An. daciae*, and hybrids demonstrated significant differences in frequencies of chromosomal inversions.

In addition to the differences identified in the frequencies of chromosomal variants between the species, our analysis also revealed distinctions between individual mosquito populations within each species (Fig. S1). For example, the gradual increase in polymorphism in the *An. messeae* 2R chromosome from south to north was observed. In southern populations all mosquitoes carried the standard variant of chromosome 2R0, but with increasing latitude, the frequencies of the standard karyotype decreased and in some northern populations, such as Surgut in the north of Western Siberia and Palevitsy in the north of Europe, this variant was represented only in a small number of heterozygous individuals. In contrast, the inverted karyotype was prevalent in the north with a frequency of more than 90%. This result suggests possible involvement of this chromosomal inversion in adaptation to cold climatic conditions. Similarly, the gradual decrease in the diversity in chromosome 3 from east to west was determined. The portions of standard karyotypes 3R0 and 3L0 steadily decreased as one goes eastward. A similar, but less pronounced, pattern of these inversion frequencies was found in *An. daciae*. Endemic inversions X2, X3, and X4 were found in Western Siberia, Latvia, and the Moscow region, respectively. In addition, frequencies of the major inversions varied greatly among the geographical regions (Fig. S1) suggesting their role in ecological adaptations of local populations or that it was a genetic drift.

To statistically identify the relationship between the frequencies of inversions and the geographical location of the mosquito collections, we constructed a correlation matrix (Table 2). A significant dependence was found for the latitude distribution of inversions on the 2R chromosome in *An. messeae*. The longitude dependence of inversion polymorphism was found for both arms of chromosome 3. The presence of such longitude and latitude clines in the inversion frequencies suggests their involvement in climatic adaptations of the mosquitoes or, alternatively, can be explained by genetic drift and dispersal from glacial refugia.

**Table 2.** Clinal variability of inversion polymorphisms in *Anopheles messeae* and *Anopheles daciae*. P-values < 0.05 or < 0.001 indicated by one or two asterisks, respectively.

|  | Latitude |  | Longitude |  |
| --- | --- | --- | --- | --- |
|  | *An. messeae* | *An. daciae* | *An. messeae* | *An. daciae* |
| X | −0.18** | −0.03 | −0.01 | −0.14** |
| 2R | −0.35** | −0.09* | 0.27** | −0.03 |
| 3R | 0.00 | −0.10* | −0.55** | −0.25** |
| 3L | −0.01 | 0.09* | −0.50** | −0.13** |

### Genetic differentiation and population structure of An. messeae and An. daciae

We estimated Hardy-Weinberg equilibrium within *An. messeae*, *An. daciae*, and their hybrids separately. The statistical analysis showed that the frequencies of homo- and heterokaryotypes, in general, did not deviate from the Hardy-Weinberg equilibrium in both species suggesting random mating within the populations (Table S4). The exceptions were found in Petergof for the 2R and 3R inversions in the *An. messeae* population (deficiency of heterozygotes in both chromosomal arms), in Solnechnogorsk for the 2R inversion in the *An. daciae* population (absence of heterozygotes), and in hybrids for the 3R inversion from Berdsk (a significant excess of heterozygotes).

To clarify the genetic differentiation between *An. messeae* and *An. daciae*, the Hendrik’s G’st indices were calculated (Table 3). Significant differentiations of 0.330 and 0.349 were found between *An. messeae* and *An. daciae* and between *An. daciae* and hybrids, respectively. At the same time, the differentiation within *An. messeae* and between *An. messeae* and hybrids were moderate (0.140). Chromosomal arms X, 2R, and 3R contributed the most to interspecies differentiation (Fst were 0.409, 0.308, and 0.492, respectively) but the difference for the 3L arm was modest (0.128). Hybrids differed moderately from both species in the X chromosomes (<1.5). The 2R arm was the main contributor to the difference between *An. messeae* and hybrids (0.301) but not between *An. daciae* and hybrids. In contrast, chromosome arms 3R and 3L differed greatly in *An. daciae* and hybrids (0.639 and 0.430) but not between *An. messeae* and hybrids (0.033 and 0.162).

**Table 3.** Genetic differentiation (Hedrick’s G’st) between *Anopheles messeae, Anopheles daciae*, and hybrids. Significant difference (p-values < 0.05) indicated by asterisks.

|  | *An. daciae*/*An. messeae* | *An. daciae*/hybrids | *An. messeae*/hybrids |
| --- | --- | --- | --- |
| X | 0.409* | 0.143* | 0.128* |
| 2R | 0.308* | 0 | 0.301* |
| 3R | 0.492* | 0.639* | 0.033* |
| 3L | 0.128* | 0.430* | 0.162* |
| Total | 0.330* | 0.349* | 0.140* |

A simple statistics analysis estimated using different approaches clearly demonstrated a higher genetic diversity of *An. messeae* and hybrids than of *An. daciae* (Table 4). For example, Menhinick indices were 3.23 in *An. messeae*, 4.04 in hybrids, but only 1.12 in *An. daciae*. Simpson indices were 0.96, 0.97, and 0.85 and Shannon indices were 3.65, 3.65, and 2.20 in *An. messeae*, hybrids, and *An. daciae*, respectively. The AMOVA test revealed a clear population structure in *An. messeae* but almost the complete absence of a population structure in *An. daciae* (Table 5). The portion of dispersion at the intrapopulation level in *An. messeae* was calculated as 61.23% (p-value < 0.001) of the total species variance, while in *An. daciae* it was only 88.49% (p-value < 0.001). In addition, we performed the isolation by distance test to estimate the genetic isolation between populations. To assess the degree of isolation by distance, the correlation coefficient between the pairwise Fst (Table S5) and the pairwise distance between the populations were calculated. To exclude the influence of the mismatch of ranges, only locations where both species were present were included in the analysis (Fig. 1). The degree of differentiation between the populations of *An. messeae* strongly depended on the distance between them (r = 0.808, p <0.001) (Fig. 5). At the same time, this dependence among *An. daciae* was more than two times weaker (r = 0.381, p <0.001). These results also suggest the presence of a clear population structure with a limited gene flow between individual populations in *An. messeae* but an absence of such population structure in *An. daciae*. PCA and hierarchical clustering based on the frequency of autosomal inversions in samples were performed to reliably separate the two species (Fig. 6). The sum of PC1 and PC2 was 86.9% of the total variance (Fig. 6A). This analysis suggests that populations of *An. messeae* do not form one distinct group. At the same time, *An. daciae* populations were grouped into one compact cluster. Thus, we determined significant differences between species at both the level of individual chromosomes and karyotypes and at the population level.

**Fig. 5.**
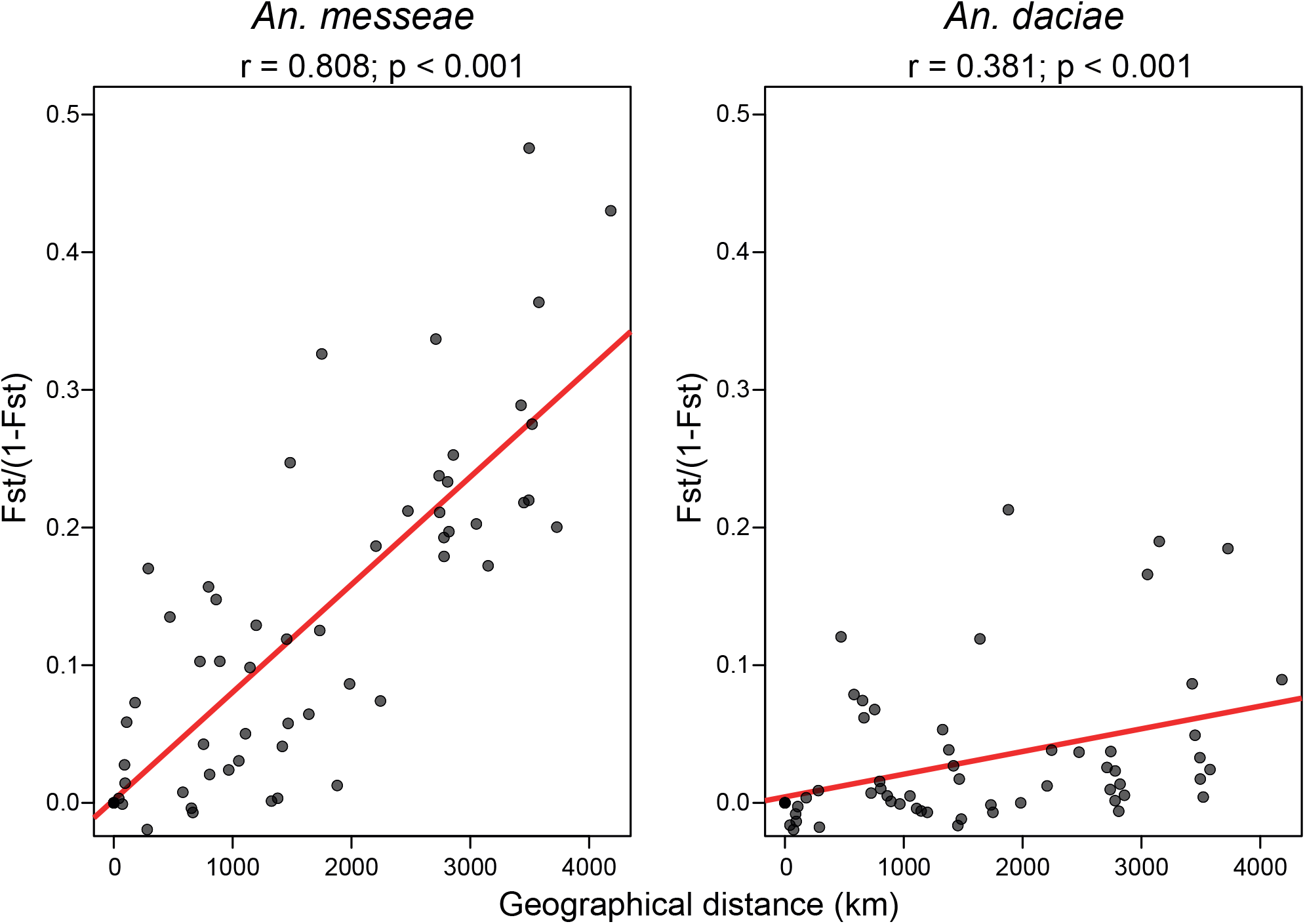
Isolation by distance within populations of *Anopheles messeae* and *Anopheles daciae*.The graph of the linear regression model is shown in red, r stands for the correlation coefficient between the pairwise Fst and the pairwise distance between populations. The X-axis shows the distance between populations in kilometers. The Y-axis indicates Fst/(1-Fst) parameter. The degree of differentiation between populations of *An.messeae* strongly depends on the distance between them, while this dependence is much weaker among *An. daciae* populations.

**Fig. 6.**
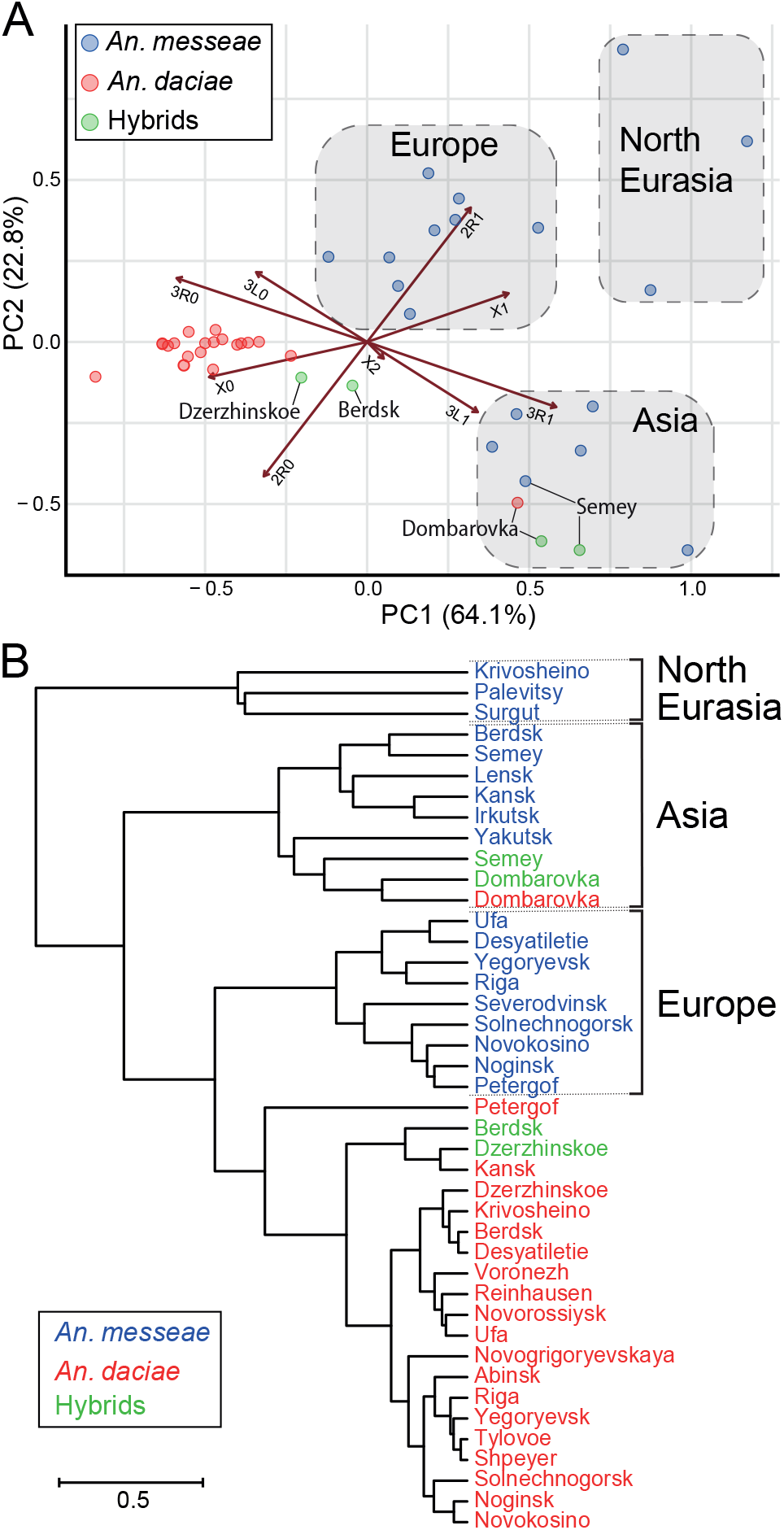
Interpopulation PCA plot (A) and hierarchical clustering dendrogram (B) based on the frequencies of the chromosomal inversions in *Anopheles messeae, Anopheles daciae*, and their hybrids. Species are indicated by different colors. Arrows on panel A indicate coordinate axes in space of frequencies of inversions X0, X1, X2, 2R0, 2R1, 3R0, 3R1, 3L0, and 3L1. Positions of populations from four locations Dzerzhinskoe, Berdsk, Semey, and Dombarovka, where most of the hybrids were found, are indicated. Three major clades of *An. mes-seae* – North Eurasian, European, and Asian – are shown by light gray color in panel A and by brackets in panel B. Based on statistical analysis, *An. messeae* is subdivided into three major clades, whereas *An. daciae* is represented by only a single population.

**Table 4.** Genetic diversity in *Anopheles messeae, Anopheles daciae*, and hybrids..

| Species | Number of sampled individuals | Number of multilocus genotypes | Menkhinik index | Corrected Simpson index ($\lambda \cdot n / n-1$) | Shannon-Wiener index |
| --- | --- | --- | --- | --- | --- |
| *An. messeae* | 830 | 93 | 3.23 | 0.96 | 3.65 |
| *An. daciae* | 923 | 34 | 1.12 | 0.85 | 2.20 |
| Hybrid | 179 | 54 | 4.04 | 0.97 | 3.65 |

**Table 5.**
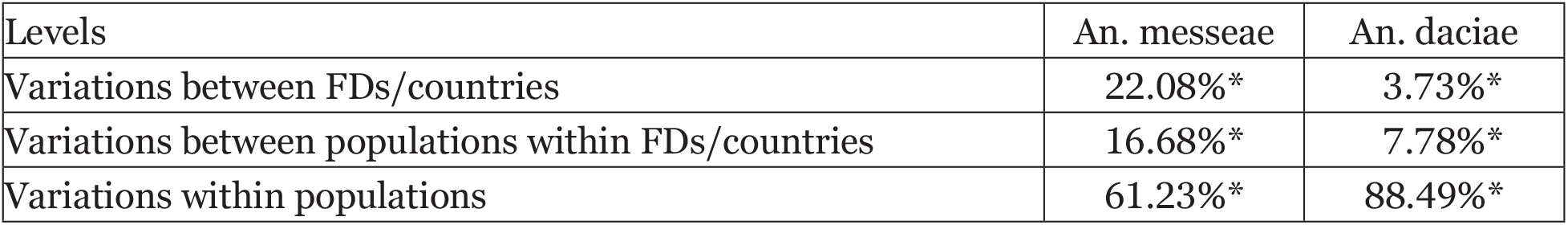
Assessment of genetic diversity and population differentiation in *Anopheles messeae* and *Anopheles daciae* based on Analysis of Molecular Variance. Significant difference (p-values < 0.05) indicated by asterisks.

When analyzing paired Fst, most of the *An. messeae* populations differed from each other, suggesting restriction of gene flow between them (Table S5). Interestingly, the two most northern populations from Europe (Palevitsy) and Asia (Surgut), which are 1200 km away from each other, were slightly different from each other but significantly distinct from the rest of the *An. messeae* populations. The same pattern is shown by the cluster dendrogramm in Fig. 6B. Thus, *An. messeae* was subdivided into three clusters, united by geographical location: Northern Eurasian, Asian, and European. A different picture was observed for *An. daciae*. Analysis of paired Fst within the species (Table S5) showed insignificant differences between populations throughout its range and, as mentioned above, *An. daciae* were grouped into one compact cluster in PCA and hierarchical clustering (Fig. 6). The exception was the population from Dombarovka (Orenburg region), which differed sharply from the rest (Fst > 0.25) and did not fall into the cluster in the PCA. At the same time, this population was included in the Asian branch of *An. messeae* in the dendrogram.

In addition to clusters of *An. messeae* and *An. daciae*, our study revealed two groups of hybrids from different locations. The two groups, southern Eurasian (Semey/Dombarovka) and Asian (Berdsk/ Tomsk), were practically identical within the group, but at the same time, were very different from each other (Fig. 6). Interestingly, the Semey/Dombarovka group was clustered with the Asian populations of *An. messeae*, whereas the Berdsk/Dzerzhinskoe group was clustered with *An. daciae*.

## Discussion

### The role of the chromosomal inversions in mosquito evolution

Chromosomal inversions represent important drivers of evolution in living organisms (Krimbas & Powell, 1992). When inversion occurs, a piece of the chromosome flips 180 degrees and produces a reverse order of the genetic material. As a result, this part of the genome becomes protected from recombination during meiosis. By capturing different sets of alleles, inversions have effects on the ecological, behavioral, and physiological adaptations of species to the natural environment (F. J. Ayala & Coluzzi, 2005). Polymorphic inversions have been shown to be responsible for epidemiologically important phenotypes in *Anopheles* populations (Coluzzi et al., 2002; Gray, Rocca, Costantini, & Besansky, 2009; Lanzaro et al., 1998; Simard et al., 2009). Reduced recombination in heterozygote inversions may promote ecological divergence leading to reproductive isolation and speciation (Hoffmann, Sgrò, & Weeks, 2004; Rieseberg, 2001). In this study, we attempted to better understand the role of chromosomal inversions in the evolution of two sibling species from the Maculipennis group, *An. messeae* and *An. daciae*. For the first time, chromosomal inversions were analyzed separately in the two species, which were identified using the ITS2 marker for species diagnostics. In total, frequencies of 11 polymorphic chromosomal inversions were evaluated in 1932 specimens from 28 locations across Eurasia. The most indicative differences between *An. messeae* and *An. daciae* were found among the inversions located on the X sex chromosome (Fig 3.). The standard arrangement X0 was present with an extremely low frequency (2%) or was almost absent in the *An. messeae* populations, while the inverted arrangement X1 was found in very high frequency (94%) in the populations of this species. In most cases the X0 arrangement was found as hetero-karyotype X01 in females and X0 hemikaryotype in males (Table 1). In contrast, both standard X0 and inverted X1 arrangements were found in almost equal frequencies of 48% and 52%, respectively, along the *An. daciae* distribution. The frequencies of other inversions were significantly less abundant than X1 inversions in the *An. messeae* populations and almost absent in the *An. daciae* populations. The endemic Siberian X2 inversion and the rare X4 inversion were only found in *An. messeae* populations with one exception where a highly abundant X2 inversion was also present in hybrids among the species in Siberia. The Latvian endemic X3 inversion (Fig. S1) was found in low frequencies in both species (<1%), and was present in *An. messeae* only in males, while in *An. daciae* it was only found in heterozygous females. Interestingly, all chromosomal inversions found in *An. messeae* were formed based on the inverted X11 arrangement, suggesting that this karyotype is ancestral for *An. messeae* (Fig. 3). Thus, we conclude that arrangement X1 was originally fixed in *An. messeae*, whereas the endemic or rare inversions on the X chromosome originated later after the two species split. In a similar situation, where one of the sibling mosquito species had a fixed inversion arrangement, but the inversion was polymorphic in another species, was found among species from the *An. gambiae* complex. In this complex, seven species were differentiated by 10 fixed chromosomal inversions (Coluzzi et al., 2002). Similar to the X1 inversion in *An. messeae* and *An. daciae*, an autosomal inversion 2La is highly polymorphic in *An. gambiae* but is fixed in its sister taxa *An. arabiensis* and *An. merus*. Detailed analyses demonstrated the ancestral state of the 2La inverted arrangement for this species complex (Kamali, Xia, Tu, & Sharakhov, 2012; Sharakhov et al., 2006). Interestingly, five fixed chromosomal inversions located on the X chromosome in the *An. gambiae* complex have no polymorphic variants in any species within the complex (Coluzzi et al., 2002). Although, five fixed chromosomal inversions have been identified among the Palearctic species of the Maculipennis group (Stegniy, 1991), five additional chromosomal inversions described in this study were polymorphic in *An. messeae* and *An. daciae* on the X chromosome.

All the wide-spread autosomal chromosomal inversions identified by this study in natural populations of *An. messeae* and *An. daciae* were present in both species indicating that their polymorphism is ancestral for both species (Fig. 3). However, the frequencies of the inverted variants 2R1, 3R1, and 3L1 were significantly higher in *An. messeae* than in *An. daciae* (0.26 versus <0.01, 0.45 versus 0.06, 0.17 versus 0.03, respectively). The rare inversions 2R4 and 2R3 were found only in *An. daciae* from Riga and in the Moscow region (Yegoryevsk), respectively, and the 3R2 and 3L4 inversions were found in the *An. messeae* northern populations of Surgut and Severodvinsk, respectively (Fig. S1). The presence of endemic chromosomal inversions suggests their involvement in local adaptations of mosquitoes. Supporting previous observations (Stegniy, Kabanova, Novikov, et al., 1976), a latitude cline was determined for 2R1 inversions with higher frequencies of standard arrangements in southern populations (Fig. S1). Longitude clines were determined for the inversions in the 3R and 3L arms. Clines were more pronounced in populations of *An. messeae* than in *An. daciae*. Chromosomal inversions were shown to be involved in adaptations in other species of mosquitoes. For example, chromosomal inversions contribute to behavioral differentiation within *An. arabiensis* (Main et al., 2016; Petrarca & Beier, 1992). The 2Rb inversion polymorphism in *An. stephensi* correlates with circadian flight activity (Jones, 1974), time of adult emergence (Coluzzi, 1972), and egg size (Coluzzi, Di Deco, & Cancrini, 1973). Intensive studies of inversion polymorphism within *An. gambiae* identified five chromosomal forms related to the inversions in the 2R arm: Bamako, Savanna, Mopti, Forest, and Bissau (Touré, 1989; Touré et al., 1994). Interestingly, these chromosomal forms may coexist in certain areas and can also replace each other geographically or seasonally (Fontenille & Simard, 2004). Gradual variations of the inversion frequencies along environmental clines were shown for inversions in *Drosophila* (Kapun, Fabian, Goudet, & Flatt, 2016; Kolaczkow-ski, Kern, Holloway, & Begun, 2011). In *An. gambiae*, the 2La inversion frequencies also change depending on the aridity gradient of the environment (D. Ayala et al., 2017, 2019; Cheng et al., 2012).

### Evolutionary histories of An. messeae and An. daciae

A better understanding of the evolutionary history of *An. messeae* and *An. daciae* in Eurasia was another important goal of this study. Here, we used ITS2 sequences and polymorphic chromosomal inversions as markers to evaluate geographical distribution, population structure, and evolutionary history of these species. Our study clearly demonstrated that *An. messeae* and *An. daciae* have different geographical distributions in Eurasia identifying them as northeastern and southwestern species, respectively (Fig. 1). The PCA and hierarchical cluster analysis, conducted based on the frequencies of the chromosomal inversions, separated *An. messeae* and *An. daciae* into two clusters (Fig. 6). In addition, our study demonstrated higher genetic diversity in *An. messeae* than *An. daciae*, suggesting the presence of a complex population structure with limited gene flow among populations of *An. messeae*. Three major clusters were identified within populations of these species: Northern Eurasian, European, and Asian. In contrast, populations of *An. daciae* clustered all together demonstrating no distinct population structure among locations in Eurasia. According to our estimation, the split between *An. messeae* and *An. daciae* occurred ~2 Mya (Yurchenko et al. 2022) and overlapped with the glaciation period (Torsvik & Cocks, 2017). The glaciation event can result in a long period of disruption of an original population of the ancestral species into different southern refugia whereas one population of mosquitoes was isolated in the Asian location and another in the European location. Over time, the two species accumulated genetic differences, which are especially pronounced in the chromosomal inversions and overall differentiation in the X chromosome (Naumenko et al., 2020). Later, when the climate became warmer, distribution of the species overlapped again but they became partially incompatible because they acquired postzygotic or prezygotic isolation barriers. Such allopatric speciation is considered a classical form of speciation commonly present in nature (Mayr, 1963). In fact, the differences in acoustic signaling of adult mosquitoes were observed between mosquitoes with different chromosomal karyotypes (Perevozkin, Printseva, Maslennikov, & Bondarchuk, 2012), which were potentially associated with *An. messeae* and *An. daciae*. Such differences may contribute to the reproductive isolation between the species. Multiple ice age cycles lasting over a thousand years became extremely severe during the last 2.4 million years (Torsvik & Cocks, 2017) and could have led to further population differentiation within *An. messeae*. Such glaciation events might have resulted in repeated restrictions of the geographical species distribution to the southern refugia and further postglacial expansion and recolonization of the northern territories. Such recolonization by a restricted number of founders reduced the allelic diversity and produced patches of genotypic homogeneity within the species along Eurasia. Based on the geographical distribution patterns and genetic diversity of the species in Eurasia, we hypothesize that *An. messeae* populated Eastern Eurasia earlier, whereas *An. daciae* colonized the Asian part of the continent more recently. As a result, *An. messeae* has a more complex population structure and higher level of inversion polymorphism in Eurasia than *An. daciae* has. We think that the appearance of *An. daciae* in Asia is a result of rapid expansion of this species to this territory due to recent climate change (Torsvik & Cocks, 2017). Fluctuations in frequencies in the X0 chromosomal variant (Stegniy et al., 2016), which is associated with *An. daciae*, indicate dramatic dynamics of the redistribution of the species in Eurasia during the last 40 years. In Western Siberia (Tomsk), frequencies of the X0 chromosome variant increased from 5 to 55% from 1974 to 2013. In Eastern Siberia (Krasnoyarsk), in 1974, the X0 variant was absent (0%) but in 2008 the frequency of this variant increased to 30%. At the same time, a tendency toward the increase in the frequency of the X0 chromosome variant was observed in a northern location (Syktyvkar). In southern Asia (Almaty), the X0 chromosome variant, which in 1974–1977 was absent (0%), had reached a frequency of 70% by 2014. In contrast, in Europe (Moscow) the X0 frequency declined from 50% (1975) to 10%. These data suggest a recent dramatic expansion of *An. daciae* in the northern, southern, and eastern territories of Eurasia with some westward expansion of *An. messeae* in Europe.

The role of the climate oscillations in speciation and population genetic processes of European and North American species was highlighted in several reviews (Emerson & Hewitt, 2005; Hewitt, 2001; Hewitt, 2004). The influence of glaciation on species evolution has been shown for mammals (Berggren, Ellegren, Hewitt, & Seddon, 2005), fish (Aguilar et al., 2019), insects (Baird et al. 2021), and plants (Tzedakis, Emerson, & Hewitt, 2013). Among mosquitoes, the presence of two groups of *An. claviger* in France, which represent unclear genetic entities, was considered as allopatric divergence in southern refugia during the last glaciation period (Schaffner, Marquine, Pasteur, & Raymond, 2003). Interestingly, the influence of glaciation on the population structure was suggested even for the tropical brackish water malaria mosquito *An. melas* (Deitz et al., 2012). Compared with its sibling species *An. gambiae, An. melas* has a higher level of genetic differentiation between the populations associated with its patchy geographical distribution, which are restricted to fragmented larval habitats in mangroves and salt marsh grass. However, the most isolated population of *An. melas* was found in Bioko Island. This island was connected to the mainland in the past but became isolated when sea levels rose after the last glaciation period 10,000-11,000 years ago. In another study, data from the microsatellite analysis of populations of the West Nile vector mosquito *Culex tarsalis* suggest that this species underwent a range expansion across the western United States within the last 375,000-560,000 years, which may have been associated with the Pleistocene glaciation events that occurred in the Midwestern and Western United States between 350,000 and 1 MYA (Venkatesan, Westbrook, Hauer, & Rasgon, 2007).

In addition to the differences in population structure between *An. messeae* and *An. daciae*, we found high variations in the frequencies of their hybrids among the locations in areas where distributions of the two species were overlapped (Fig. 1). The frequencies of hybrids were very low in northern Europe from 0% (Severodvinsk) to 1.59 % (Solnechnogorsk) but higher, up to 7%, in southern Europe (Voronezh). In contrast, in two locations in southwestern Siberia (Berdsk and Dzerzhinskoe), one Southern Ural location (Dombarovka), and one Kazakhstan location (Semey), the hybrid frequencies were very high (49.43%, 8.89%, 78.13%, and 54.26%, respectively). Surprisingly, only *An. messeae*, in addition to the hybrids, was present in the Semey location and only *An. daciae*, in addition to the hybrids, was present in the Dombarovka location. These data could have multiple interpretations from just the simple presence of recombinants in rDNA sequences in one or in both species in some geographical areas to an existence of an additional cryptic taxa in Central Asia that is ancestral to both *An. messeae* and *An. daciae*. Southern Eurasia can also be considered as a hybrid zone between the two species. Interestingly, low frequencies of the heterozygote X01, which may represent potential hybrids between the two species, were always observed in the western part of the species distribution but were interpreted as disruptive selection that favors the X00 and X11 homozygotes at different times of the breeding and wintering seasons (Stegniy et al., 2016).

Hybridization between closely related species has been shown to be variable in other mosquitoes. For example, the number of hybrids between two incipient species, *An. gambiae* and *An. coluzzii*, from the *An. gambiae* complex varies among the locations from 0.2% in Coastal areas of Africa to 24% in Guinea-Bissau (Oliveira et al., 2008) and even up to 42% in some surrounding areas (Nwakanma et al., 2013). More recent genomic analyses conducted based on 1142 individual mosquitoes from 13 African countries suggest the presence of three major groupings among the individuals: 1) *An. gambiae* from west, central, and near-east Africa, 2) *An. coluzzii* from west Africa, and 3) individuals with uncertain species status from far-west Africa (Anopheles gambiae 1000 Genomes Consortium, 2020). Patterns of interspecies hybridization vary in *Culex* mosquitoes. For example, massive hybrid zones were found between mosquitoes from the *Culex pipiens* complex. Diapausing in winter *Cx. pipiens* and non-diapausing *Cx. quinquefasciatus* form large hybrid zones where their distribution overlaps in North America, Africa, and Asia (Aarde-ma, Olatunji, & Fonseca, 2022; Farajollahi, Fonseca, Kramer, & Marm Kilpatrick, 2011). Two eco-physiological forms or subspecies, *Cx. p. pipiens* and *Cx. p. molestus*, which are behaviorally divergent and are truly isolated in northern Europe, but are gradually becoming well-mixed, intermediate populations in North Africa (Haba & McBride, 2022). A number of studies suggest that *Cx. p. pipiens* and *Cx. p. molestus* represent different evolutionary entities (Fonseca et al., 2004; Gomes et al., 2015; Yurchenko et al., 2020).

We think that additional whole-genome sequencing approach of multiple individuals of *An. messeae*, *An. daciae*, and their hybrids from different locations in Eurasia has to be considered for the future studies. Such analysis will help for better understanding population structure, evolutionary history of these two sibling species in Eurasia, and the nature of hybrids between them.

## Conclusions

Chromosomal inversions represent powerful markers for better understanding of genetic mechanisms of evolution and phylogeography of mosquitoes. Analysis of the chromosomal inversions conducted in this study indicated their role in the evolution of the two sibling species of malaria mosquitoes *An. messeae* and *An. daciae* in Eurasia. Existence of species-specific inversions in the X chromosome of these species implies their potential involvement in speciation. The presence of longitude and latitude clines in Eurasia associated with frequencies of the autosomal inversions suggests their role in adaptation to different climate conditions. We demonstrated that *An. messeae* and *An. daciae* differ from each other by their geographical distribution, population structure, and evolutionary history in Eurasia. The study showed that *An. messeae* has much higher diversity and a more complex population structure, while *An. daciae* has low diversity and a more homogeneous structure of its populations. We hypothesize that glaciation events in Eurasia may have contributed to the original split between *An. messeae* and *An. daciae* and explain the higher genetic diversity within *An. messeae*. Our data suggest that *An. messeae* populated Eastern Eurasia earlier, whereas *An. daciae* spread in Asia more recently. Detailed knowledge of the population structure and evolutionary history of the important malaria vectors *An. messeae* and *An. daciae* will help to predict the patterns of malaria transmission in Eurasia.

## Declarations

### Ethics approval and consent to participate

Not applicable.

### Consent for publication

Not applicable.

### Availability of data and material

All the data are available in the text, figures, and tables of this article. Consensus ITS2 sequences of all variants found in this study are available in GenBank under the accession numbers shown in Figure 2.

### Authors’ contribution

Conceptualization, M.V.S., I.I.B., and M.I.G.; methodology, I.I.B., D.A.K., A.A.Y., and A.V.M.; investigation, I.I.B., D.A.K., J.M.H., A.V.M., and M.I.G.; writing—original draft preparation, M.V.S., I.I.B. and D.A.K.; writing—review and editing, M.V.S., I.I.B., E.N.B., I.V.S., and D.A.K.; supervision, M.V.S.; project administration, E.M.B.; funding acquisition, M.V.S., E.M.B., and M.I.G.; mosquito collections, I.I.B., A.V.M., M.I.G., V.A.B., G.N.A., A.K.S., N.B., I.V.S., M.V.S. All authors have read and agreed to the published version of the manuscript.

## Acknowledgements

We thank the Russian Science Foundation for funding the ITS2 sequencing (grant № 19-14-00130 to M.V.S.) and for supporting the inversion polymorphism analysis (grant 22-24-00183 to M.I.G). We also thank the Institute of Cytology and Genetics for funding the statistical analysis (FWNR-2022-0015 budget project) and Janet Webster for editing the text.

## Supplementary Materials

**Fig. S1.**
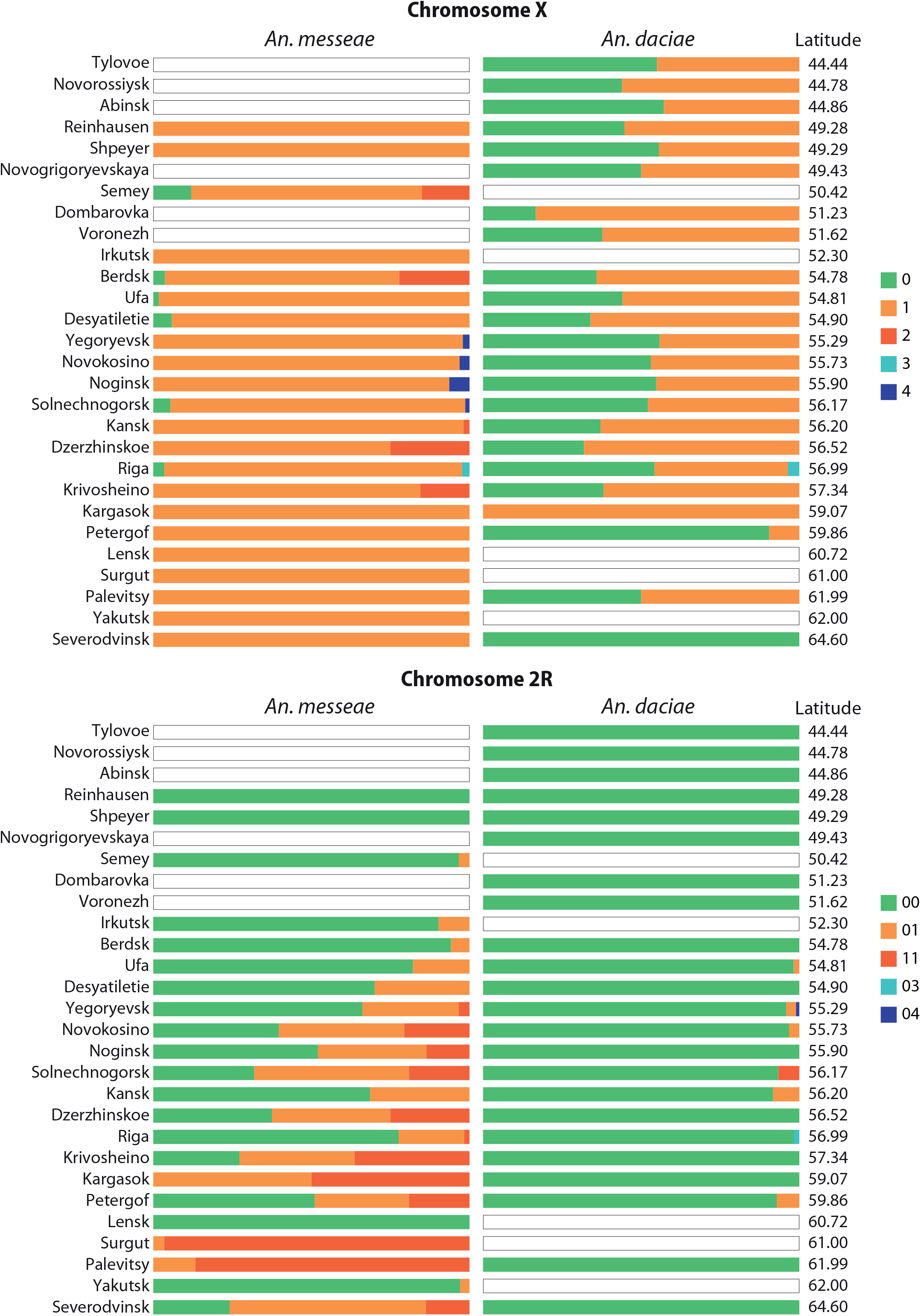
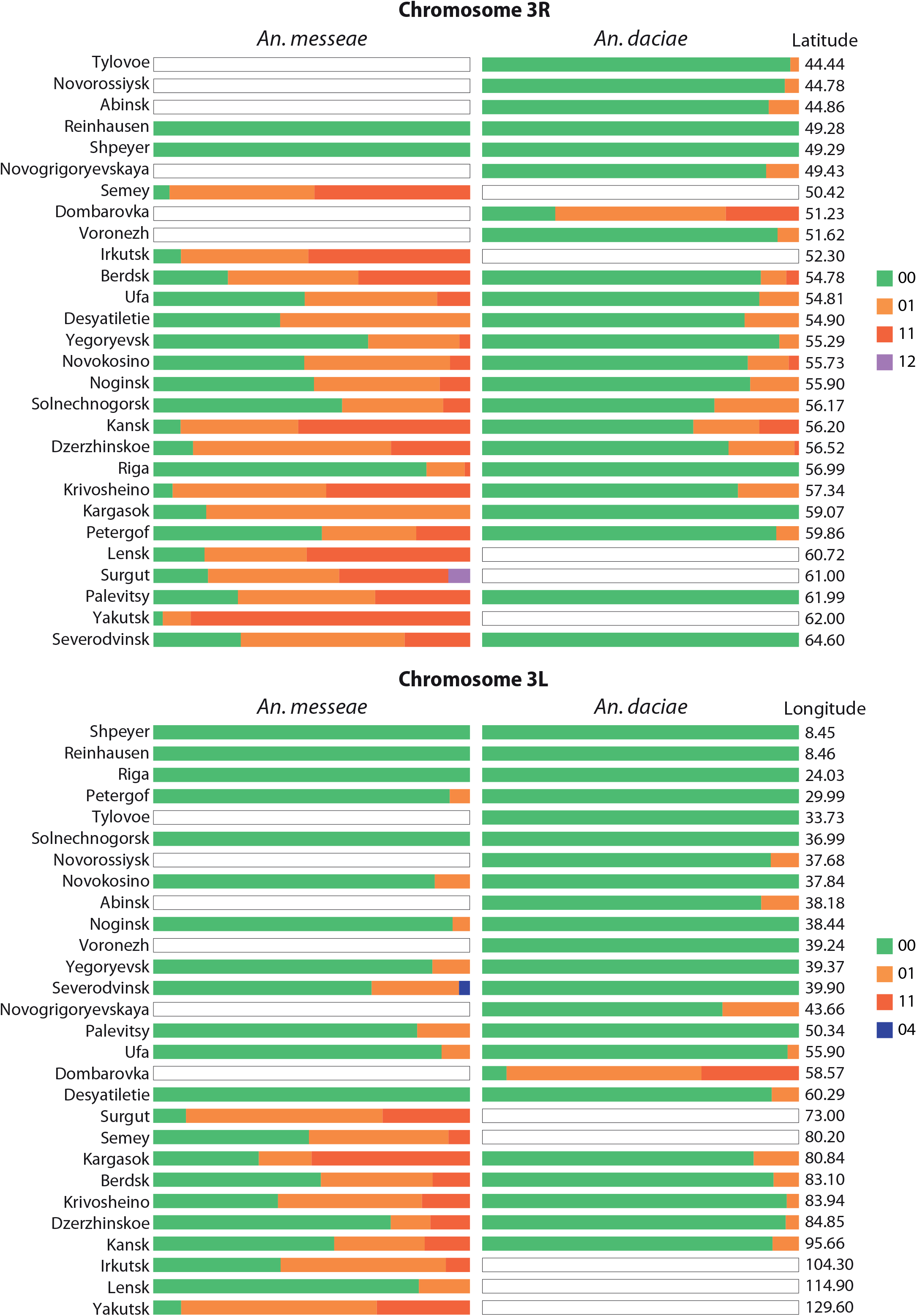
The frequencies of chromosomal inversions in *Anopheles messeae*, *Anopheles daciae*, and hybrids in studied populations. Proportions of standard, inverted, and heterozygote arrangements are shown by different colors. Locations of the collection sites are represented in ascending order of longitude for X and 2R chromosomes and in ascending order of latitude for chromosomes 3R and 3L. In both cases, one can see a gradual change in the inversion frequencies depending on coordinates.

**Table S1.** Locations, dates, and counts of the mosquitoes in the sample collections.

| Country and Federal District | Region | Location | Coordinates | Year | Count |
| --- | --- | --- | --- | --- | --- |
| Russia, Central FD | Moskovskaya Oblast | Noginsk | 55.89927 38.438304 | 2017 | 24 |
|  |  |  |  | 2016 | 99 |
|  | Moscow Federal City | Novokosino | 55.73406 37.842629 | 2016 | 94 |
|  | Moskovskaya Oblast | Solnechnogorsk | 56.17424 36.985894 | 2019 | 120 |
|  |  | Yegoryevsk | 55.28831 39.367628 | 2017 | 258 |
|  |  |  |  | 2016 | 87 |
|  | Voronezhskaya Oblast | Voronezh | 51.62370 39.238098 | 2018 | 110 |
| Russia, Southern FD | Sevastopol' Federal City | Tylovoe | 44.44174 33.727469 | 2017 | 100 |
|  |  |  |  | 2019 | 152 |
|  | Krasnodyarskiy Kray | Abinsk | 44.86238 38.183556 | 2018 | 109 |
|  |  | Novorossiysk | 44.78080 37.680039 | 2018 | 110 |
|  | Volgogradskaya Oblast | Novogrigor'yevskaya | 49.43001 43.662200 | 2020 | 58 |
| Russia, Volga FD | Republic of Bashkortostan | Ufa | 54.80943 55.899455 | 2017 | 209 |
|  | Orenburgskaya Oblast | Dombarovka | 51.22609 58.567493 | 2020 | 119 |
| Russia, Northwestern FD | Arkhangel'skaya Oblast | Severodvinsk | 64.60000 39.900000 | 2020 | 118 |
|  | Komi Republic | Palevitsy | 61.98569 50.336472 | 2019 | 118 |
|  | Sankt-Peterburg Federal City | Petergof | 59.86241 29.992752 | 2019 | 118 |
| Russia, Ural FD | Chelyabinskaya Oblast | Desyatiletie | 54.89772 60.289069 | 2019 | 114 |
|  | Khanty-Mansiyskiy Autonomous Okrug | Surgut | 61.00000 73.000000 | 2019 | 120 |
| Russia, Siberian FD | Irkutskaya Oblast | Irkutsk | 52.30077 104.346800 | 2017 | 97 |
|  | Krasnoyarskiy Kray | Kansk | 56.20486 95.663929 | 2019 | 119 |
|  | Novosibirskaya Oblast | Berdsk | 54.77658 83.098135 | 2017 | 99 |
|  | Tomskaya Oblast | Dzerzhinskoe | 56.51615 84.853488 | 2017 | 147 |
|  |  | Kargasok | 59.06622 80.843556 | 2018 | 110 |
|  |  | Krivosheino | 57.34198 83.941120 | 2019 | 89 |
| Russia, Far Eastern FD | Republic of Sakha (Yakutia) | Lensk | 60.72181 114.877915 | 2019 | 120 |
|  |  | Yakutsk | 62.00219 129.627405 | 2019 | 102 |
| Kazakhstan | East Kazakhstan (Shyghys Qazaqstan) | Semey | 50.41750 80.202778 | 2019 | 118 |
|  |  |  |  | 2018 | 100 |
| Germany | Baden-Württemberg | Reinhausen | 49.27667 8.460436 | 2018 | 120 |
|  | Rheinland-Pfalz | Speyer | 49.29257 8.453797 | 2018 | 110 |
| Latvia | Riga | Riga | 56.98661 24.032325 | 2018 | 120 |

Table S2. Metadata for the samples used for the statistical analyses (data not shown).

**Table S3.** Distribution of *Anopheles messeae, Anopheles daciae*, and their hybrids in Eurasia.

| Location | *An.daciae* | *An.messeae* | Hybrid |
| --- | --- | --- | --- |
| Noginsk | 34 (27.87%) | 87 (71.31%) | 1 (0.82%) |
| Novokosino | 31 (32.99%) | 63 (67.02%) |  |
| Solnechnogorsk | 15 (23.81%) | 47 (74.60%) | 1 (1.59%) |
| Yegoryevsk | 146 (54.89%) | 118 (44.36%) | 2 (0.75%) |
| Voronezh | 93 (93.00%) |  | 7 (7%) |
| Tylovoe | 39 (100.0%) |  |  |
| Abinsk | 85 (100.0%) |  |  |
| Novorossiysk | 45 (100.0%) |  |  |
| Novogrigoryevskaya | 28 (100.0%) |  |  |
| Ufa | 84 (41.18%) | 119 (58.33%) | 1 (0.49%) |
| Dombarovka | 13 (20.31%) | 1 (1.56%) | 50 (78.13%) |
| Severodvinsk | 6 (17.14%) | 29 (82.86%) |  |
| Palevitsy | 1 (3.13%) | 30 (93.75%) | 1 (3.125%) |
| Petergof | 14 (22.58%) | 48 (77.42%) |  |
| Desyatiletie | 35 (77.78%) | 10 (22.22%) |  |
| Surgut |  | 29 (100%) |  |
| Irkutsk |  | 93 (100%) |  |
| Kansk | 24 (39.34%) | 35 (57.38%) | 2 (3.28%) |
| Berdsk | 25 (28.74%) | 19 (21.84%) | 43 (49.43%) |
| Dzerzhinskoe | 112 (82.96%) | 11 (8.15%) | 12 (8.89%) |
| Kargasok | 7 (53.85%) | 6 (46.15%) | (0%) |
| Krivosheino | 27 (44.26%) | 33 (54.10%) | 1 (1.64%) |
| Lensk |  | 31 (100%) |  |
| Yakutsk |  | 34 (100%) |  |
| Semey |  | 59 (46.46%) | 68 (53.54%) |
| Reinhausen | 32 (96.97%) | 1 (3.03%) |  |
| Speyer | 105 (98.13%) | 2 (1.87%) |  |
| Riga | 57 (49.57%) | 58 (50.43%) |  |

**Table S4.** Deviation from Hardy-Weinberg equilibrium (HWE) in mosquito populations. Populations with a significant deviation from HWE are shown by asterisks.

| Locality | *An. daciae* |  |  | *An. messeae* |  |  | Hybrid |  |  |
| --- | --- | --- | --- | --- | --- | --- | --- | --- | --- |
|  | 2R | 3R | 3L | 2R | 3R | 3L | 2R | 3R | 3L |
| Noginsk |  | 1.000 |  | 0.101 | 0.778 | 1.000 |  |  |  |
| Novokosino | 1.000 | 0.233 |  | 0.191 | 0.546 | 1.000 |  |  |  |
| Solnechnogorsk | 0.034* | 1.000 |  | 1.000 | 0.426 |  |  |  |  |
| Yegoryevsk | 1.000 | 1.000 |  | 1.000 | 1.000 | 1.000 |  |  |  |
| Voronezh |  | 1.000 |  |  |  |  |  |  |  |
| Tylovoe |  | 1.000 |  |  |  |  |  |  |  |
| Abinsk |  | 1.000 | 1.000 |  |  |  |  |  |  |
| Novorossiysk |  | 1.000 | 1.000 |  |  |  |  |  |  |
| Novogrigoryevskaya |  | 1.000 | 1.000 |  |  |  |  |  |  |
| Ufa | 1.000 | 1.000 | 1.000 | 1.000 | 0.781 | 1.000 |  |  |  |
| Dombarovka |  | 1.000 | 0.566 |  |  |  |  | 0.051 | 0.768 |
| Severodvinsk |  |  |  | 0.268 | 1.000 | 1.000 |  |  |  |
| Palevitsy |  |  |  | 1.000 | 0.479 | 1.000 |  |  |  |
| Petergof | 1.000 | 1.000 |  | 0.024* | 0.041* | 1.000 |  |  |  |
| Desyatiletie |  | 1.000 | 1.000 |  |  |  |  |  |  |
| Surgut |  |  |  | 1.000 | 0.699 | 0.250 |  |  |  |
| Irkutsk |  |  |  | 1.000 | 0.804 | 0.161 |  |  |  |
| Kansk | 1.000 | 0.063 | 1.000 | 0.566 | 0.681 | 0.095 |  |  |  |
| Berdsrk |  | 0.121 | 1.000 | 1.000 | 0.624 | 0.573 | 1.000 | 0.012* | 1.000 |
| Dzerzhinskoe |  | 1.000 | 1.000 |  |  |  |  | 1.000 | 1.000 |
| Kargasok |  |  |  |  |  |  |  |  |  |
| Krivosheino |  | 1.000 | 1.000 | 0.160 | 0.680 | 1.000 |  |  |  |
| Lensk |  |  |  |  | 0.211 | 1.000 |  |  |  |
| Yakutsk |  |  |  | 1.000 | 0.146 | 0.155 |  |  |  |
| Semey |  |  |  | 1.000 | 0.516 | 0.754 |  | 0.478 | 0.452 |
| Reinhausen |  |  |  |  |  |  |  |  |  |
| Speyer |  |  |  |  |  |  |  |  |  |
| Riga |  |  |  | 1.000 | 0.285 |  |  |  |  |

**Table S5.** Paired Fst values. Significant Fst values (p<0.05) are labeled in bold.

| **Hybrids** |  |  |  |
| --- | --- | --- | --- |
|  | Dombrovka | Dzerzhinskoe | Berdsk |
| Dzerzhinskoe | **0.267** |  |  |
| Berdsk | **0.229** | -0.021 |  |
| Semey | 0.025 | **0.340** | **0.284** |

Table S5 (continuation).
| An.daciae |  |  |  |  |  |  |  |  |  |  |  |  |  |  |  |  |  |  |  |
| --- | --- | --- | --- | --- | --- | --- | --- | --- | --- | --- | --- | --- | --- | --- | --- | --- | --- | --- | --- |
|  | Dombarovka | Dzerzhinskoe | Yegoryevsk | Kansk | Krivoshino | Berdsk | Novogrigor'yevskaya | Reinhausen | Solnechnogorsk | Novorossiysk | Voronezh | Ufa | Noginsk | Novokosino | Abinsk | Tylovoe | Riga | Speyer | Petergof |
| Dzerzhinskoe | 0.410 |  |  |  |  |  |  |  |  |  |  |  |  |  |  |  |  |  |  |
| Yegoryevsk | **0.509** | 0.061 |  |  |  |  |  |  |  |  |  |  |  |  |  |  |  |  |  |
| Kansk | **0.279** | 0.007 | 0.087 |  |  |  |  |  |  |  |  |  |  |  |  |  |  |  |  |
| Krivoshino | **0.388** | -0.008 | 0.026 | 0.007 |  |  |  |  |  |  |  |  |  |  |  |  |  |  |  |
| Berdsk | **0.379** | -0.010 | 0.0373 | 0.010 | -0.018 |  |  |  |  |  |  |  |  |  |  |  |  |  |  |
| Novo-grigoryevskaya | **0.346** | 0.0301 | 0.0188 | 0.039 | 0.006 | 0.005 |  |  |  |  |  |  |  |  |  |  |  |  |  |
| Reinhausen | **0.495** | 0.022 | 0.016 | 0.068 | 0.001 | 0.002 | 0.016 |  |  |  |  |  |  |  |  |  |  |  |  |
| Solnechnogorsk | **0.353** | 0.018 | 0.004 | 0.004 | -0.006 | 0.006 | 0.007 | 0.022 |  |  |  |  |  |  |  |  |  |  |  |
| Novorossiysk | **0.448** | 0.014 | 0.021 | 0.049 | -0.005 | -0.007 | 0.001 | -0.009 | 0.016 |  |  |  |  |  |  |  |  |  |  |
| Voronezh | **0.516** | 0.011 | 0.030 | 0.057 | -0.004 | -0.005 | 0.025 | -0.007 | 0.024 | -0.004 |  |  |  |  |  |  |  |  |  |
| Ufa | **0.443** | 0.0155 | 0.005 | 0.037 | -0.007 | -0.002 | 0.003 | -0.002 | -0.007 | -0.0029 | 0.002 |  |  |  |  |  |  |  |  |
| Noginsk | **0.433** | 0.042 | -0.008 | 0.049 | 0.010 | 0.023 | 0.008 | 0.0175 | -0.014 | 0.017 | 0.027 | -0.004 |  |  |  |  |  |  |  |
| Novokosino | **0.416** | 0.030 | -0.003 | 0.033 | 0.002 | 0.014 | 0.008 | 0.015 | -0.020 | 0.014 | 0.021 | -0.006 | -0.016 |  |  |  |  |  |  |
| Abinsk | **0.437** | 0.045 | 0.001 | 0.063 | 0.014 | 0.021 | -0.003 | 0.014 | 0.003 | 0.010 | 0.026 | 0.001 | -0.006 | -0.002 |  |  |  |  |  |
| Tylovoe | **0.500** | 0.064 | -0.008 | 0.092 | 0.028 | 0.040 | 0.016 | 0.014 | 0.010 | 0.019 | 0.031 | 0.004 | -0.010 | -0.002 | -0.002 |  |  |  |  |
| Riga | **0.523** | 0.048 | -0.001 | 0.090 | 0.017 | 0.024 | 0.016 | -0.003 | 0.016 | 0.005 | 0.012 | 0.000 | 0.001 | 0.005 | 0.004 | -0.006 |  |  |  |
| Petergof | **0.495** | 0.216 | 0.068 | 0.185 | 0.166 | 0.190 | 0.106 | 0.186 | 0.079 | 0.169 | 0.207 | 0.119 | 0.062 | 0.074 | 0.072 | 0.072 | 0.121 | 0.099 |  |
| Desyatiletie | **0.388** | -0.008 | 0.053 | 0.012 | -0.012 | -0.017 | 0.015 | 0.011 | 0.017 | 0.001 | 0.003 | 0.009 | 0.039 | 0.027 | 0.035 | 0.057 | 0.038 | 0.060 | 0.213 |

Table S5 (continuation).
| An. messeae |  |  |  |  |  |  |  |  |  |  |  |  |  |  |  |  |  |
| --- | --- | --- | --- | --- | --- | --- | --- | --- | --- | --- | --- | --- | --- | --- | --- | --- | --- |
|  | Yegoryevsk | Kansk | Krivoshheino | Berdsk | Palevitsy | Severod-<br>vinsk | Solnechno-<br>gorsk | Semey | Surgut | Ufa | Noginsk | Novokosino | Irkutsk | Riga | Yakutsk | Petergof | Lensk |
| Kansk | **0.289** |  |  |  |  |  |  |  |  |  |  |  |  |  |  |  |  |
| Krivoshheino | **0.337** | 0.103 |  |  |  |  |  |  |  |  |  |  |  |  |  |  |  |
| Berdsk | 0.211 | 0.0216 | 0.170 |  |  |  |  |  |  |  |  |  |  |  |  |  |  |
| Palevitsy | **0.473** | **0.415** | 0.185 | **0.488** |  |  |  |  |  |  |  |  |  |  |  |  |  |
| Severodvinsk | 0.133 | 0.119 | 0.072 | 0.129 | 0.188 |  |  |  |  |  |  |  |  |  |  |  |  |
| Solnechnogorsk | 0.073 | **0.275** | 0.233 | **0.253** | **0.263** | 0.046 |  |  |  |  |  |  |  |  |  |  |  |
| Semey | **0.302** | 0.017 | 0.196 | 0.011 | **0.491** | 0.193 | **0.324** |  |  |  |  |  |  |  |  |  |  |
| Surgut | **0.575** | **0.422** | 0.153 | **0.488** | 0.219 | **0.308** | **0.442** | **0.500** |  |  |  |  |  |  |  |  |  |
| Ufa | 0.031 | 0.212 | **0.326** | 0.125 | **0.503** | 0.139 | 0.129 | 0.211 | **0.595** |  |  |  |  |  |  |  |  |
| Noginsk | 0.028 | 0.218 | 0.238 | 0.179 | **0.333** | 0.042 | 0.014 | **0.259** | **0.481** | 0.050 |  |  |  |  |  |  |  |
| Novokosino | 0.059 | 0.220 | 0.193 | 0.197 | **0.253** | 0.019 | -0.001 | **0.275** | **0.413** | 0.098 | 0.003 |  |  |  |  |  |  |
| Irkutsk | **0.304** | 0.005 | 0.181 | 0.010 | **0.489** | 0.192 | **0.333** | 0.007 | **0.491** | 0.220 | **0.263** | **0.277** |  |  |  |  |  |
| Riga | 0.024 | **0.430** | **0.476** | **0.364** | **0.613** | **0.272** | 0.157 | **0.411** | **0.694** | 0.086 | 0.103 | 0.148 | **0.410** |  |  |  |  |
| Yakutsk | **0.559** | 0.145 | **0.265** | 0.214 | **0.639** | **0.393** | **0.547** | 0.136 | **0.555** | **0.497** | **0.495** | **0.488** | 0.111 | **0.686** |  |  |  |
| Petergof | 0.043 | 0.200 | 0.203 | 0.172 | **0.300** | 0.023 | 0.008 | 0.249 | **0.450** | 0.064 | -0.007 | -0.004 | **0.254** | 0.135 | **0.488** |  |  |
| Lensk | **0.291** | 0.062 | **0.268** | 0.055 | **0.562** | 0.202 | **0.316** | 0.048 | **0.604** | 0.165 | 0.225 | **0.256** | 0.065 | **0.442** | **0.342** | 0.225 |  |
| Desyatiletie | 0.001 | 0.187 | 0.247 | 0.119 | **0.447** | 0.074 | 0.058 | 0.199 | **0.537** | -0.019 | 0.003 | 0.041 | 0.223 | 0.074 | **0.523** | 0.013 | 0.191 |

## References

Aardema, M. L., Olatunji, S. K., & Fonseca, D. M. (2022). The enigmatic *Culex pipiens* (Diptera: Culicidae) species complex: phylogenetic challenges and opportunities from a notoriously tricky mosquito group. Annals of the Entomological Society of America, 115(1), 95–104. doi: 10.1093/aesa/saab038

Aguilar, C., Miller, M. J., Loaiza, J. R., González, R., Krahe, R., & De León, L. F. (2019). Tempo and mode of allopatric divergence in the weakly electric fish *Sternopygus dariensis* in the Isthmus of Panama. Scientific Reports, 9(1). doi: 10.1038/S41598-019-55336-Y

Anopheles gambiae 1000 Genomes Consortium. (2020). Genome variation and population structure among 1142 mosquitoes of the African malaria vector species *Anopheles gambiae* and *Anopheles coluzzii*. Genome Research, 30(10), 1533–1546. doi: 10.1101/gr.262790.120

Artemov, G. N., Fedorova, V. S., Karagodin, D. A., Brusentsov, I. I., Baricheva, E. M., Sharakhov, I. V.,… Sharakhova, M. V. (2021). New cytogenetic photomap and molecular diagnostics for the cryptic species of the malaria mosquitoes *Anopheles messeae* and *Anopheles daciae* from Eurasia. Insects, 12(9), 835. doi: 10.3390/insects12090835

Ayala, D., Acevedo, P., Pombi, M., Dia, I., Boccolini, D., Costantini, C.,… Fontenille, D. (2017). Chromosome inversions and ecological plasticity in the main African malaria mosquitoes. Evolution; International Journal of Organic Evolution, 71(3), 686–701. doi: 10.1111/evo.13176

Ayala, D., Zhang, S., Chateau, M., Fouet, C., Morlais, I., Costantini, C.,… Besansky, N. J. (2019). Association mapping desiccation resistance within chromosomal inversions in the African malaria vector *Anopheles gambiae*. Molecular Ecology, 28(6), 1333–1342. doi: 10.1111/mec.14880

Ayala, F. J., & Coluzzi, M. (2005). Chromosome speciation: humans, Drosophila, and mosquitoes. Proceedings of the National Academy of Sciences of the United States of America, 102 Suppl, 6535–6542. doi: 10.1073/pnas.0501847102

Beklemishev, V. N. (1948). *Anopheles* species in the USSR and neighboring Asian countries, their distribution and participation in the transmission of malaria (review). Medical Parasitology and Parasitic Diseases, 17(3), 201–209.

Berggren, K. T., Ellegren, H., Hewitt, G. M., & Seddon, J. M. (2005). Understanding the phylogeographic patterns of European hedgehogs, Erinaceus concolor and E. europaeus using the MHC. Heredity (Edinb) 95(1), 84–90. doi: 10.1038/sj.hdy.6800694

Bertola, M., Mazzucato, M., Pombi, M., & Montarsi, F. (2022). Updated occurrence and bionomics of potential malaria vectors in Europe: a systematic review (2000-2021). Parasites & Vectors, 15(1), 88. doi: 10.1186/s13071-022-05204-y

Bezzhonova, O. V., & Goryacheva, I. I. (2008). Intragenomic Heterogeneity of rDNA Internal Transcribed Spacer 2 in *Anopheles messeae* (Diptera: Culicidae). Journal of Medical Entomology, 45(3), 337–341. doi: fgfge10.1603/0022-2585(2008)45[337:ihorit]2.0.co;2

Blažejová, H., Šebesta, O., Rettich, F., Mendel, J., Čabanová, V., Miterpáková, M.,… Rudolf, I. (2018). Cryptic species *Anopheles daciae* (Diptera: Culicidae) found in the Czech Republic and Slovakia. Parasitology Research, 117(1), 315–321. doi: 10.1007/s00436-017-5670-0

Burlak, V. A., & Gordeev, M. I. (1998). The effect of infection by the entomopathogenic bacterium Bacillus thuringiensis on the spread of microsporidia in an inversion-polymorphic population of the malarial mosquito *Anopheles messeae* (Diptera: Culicidae). Parazitologiia, 32(3), 264–267. Retrieved from http://www.ncbi.nlm.nih.gov/pubmed/9702802

Calzolari, M., Desiato, R., Albieri, A., Bellavia, V., Bertola, M., Bonilauri, P.,… Montarsi, F. (2021). Mosquitoes of the Maculipennis complex in Northern Italy. Scientific Reports, 11(1), 6421. doi: 10.1038/s41598-021-85442-9

Cheng, C., White, B. J., Kamdem, C., Mockaitis, K., Costantini, C., Hahn, M. W., & Besansky, N. J. (2012). Ecological genomics of *Anopheles gambiae* along a latitudinal cline: A population-resequencing approach. Genetics, 190(4), 1417–1432. doi: 10.1534/genet-ics.111.137794

Chretien, J. P., Anyamba, A., Small, J., Britch, S., Sanchez, J. L., Halbach, A. C.,… Linthicum, K. J. (2015). Global climate anomalies and potential infectious disease risks: 2014-2015. PLoS Currents, 7(OUTBREAKS), 2014–2015. doi: 10.1371/currents.outbreaks.95fbc4a8fb4695e049baabfc2fc8289f

Coluzzi, M. (1972). Inversion polymorphism and adult emergence in *Anopheles stephensi*. Science (New York, N.Y.), 176(4030), 59–60. doi: 10.1126/science.176.4030.59

Coluzzi, M., Di Deco, M. A., & Cancrini, G. (1973). Further observations on the egg length in *Anopheles stephensi* in relation to chromosomal polymorphism. Parassitologia, 15(3), 213–215. Retrieved from http://www.ncbi.nlm.nih.gov/pubmed/4807776

Coluzzi, M., Sabatini, A., Della Torre, A., Di Deco, M. A., & Petrarca, V. (2002). A poly-tene chromosome analysis of the *Anopheles gambiae* species complex. Science, 298(5597), 1415–1418. doi: 10.1126/sci-ence.1077769

Culverwell, C. L., Vapalahti, O. P., & Harbach, R. E. (2020). *Anopheles daciae*, a new country record for Finland. Medical and Veterinary Entomology, 34(2), 145–150. doi: 10.1111/mve.12431

Czajka, C., Weitzel, T., Kaiser, A., Pfitzner, W. P., & Becker, N. (2020). Species composition, geographical distribution and seasonal abundance of the *Anopheles maculipennis* complex along the Upper Rhine, Germany. Parasitology Research, 119(1), 75–84. doi: 10.1007/s00436-019-06551-z

Danabalan, R., Monaghan, M. T., Ponsonby, D. J., & Linton, Y.-M. (2014). Occurrence and host preferences of *Anopheles maculipennis* group mosquitoes in England and Wales. Medical and Veterinary Entomology, 28(2), 169–178. doi: 10.1111/mve.12023

Daskova, N. G., & Rasnicyn, S. P. (1982). Review of data on susceptibility of mosquitos in the USSR to imported strains of malaria parasites. Bulletin of the World Health Organization, 60(6), 893–897.

Deitz, K. C., Athrey, G., Reddy, M. R., Overgaard, H. J., Matias, A., Jawara, M.,… Slotman, M. A. (2012). Genetic isolation within the malaria mosquito *Anopheles melas*. Mol Ecol, 21(18), 4498–4513. doi: 10.5061/dryad.d15d4

Emerson, B. C., & Hewitt, G. M. (2005). Phylogeography. Current Biology: CB, 15(10). doi: 10.1016/J.CUB.2005.05.016

Excoffier, L., Smouse, P. E., & Quattro, J. M. (1992). Analysis of molecular variance inferred from metric distances among DNA haplotypes: application to human mitochondrial DNA restriction data. Genetics, 131(2),479–491. Retrieved from http://www.ncbi.nlm.nih.gov/pubmed/1644282

Farajollahi, A., Fonseca, D. M., Kramer, L. D., & Marm Kilpatrick, A. (2011). “Bird biting” mosquitoes and human disease: a review of the role of *Culex pipiens* complex mosquitoes in epidemiology. Infection, Genetics and Evolution: Journal of Molecular Epidemiology and Evolutionary Genetics in Infectious Diseases, 11(7), 1577–1585. doi: 10.1016/j.meegid.2011.08.013

Fonseca, D. M., Keyghobadi, N., Malcolm, C. A., Mehmet, C., Schaffner, F., Mogi, M.,… Wilkerson, R. C. (2004). Emerging vectors in the *Culex pipiens* complex. Science (New York, N.Y.), 303(5663), 1535–1538. doi: 10.1126/science.1094247

Fontenille, D., & Simard, F. (2004). Unravelling complexities in human malaria transmission dynamics in Africa through a comprehensive knowledge of vector populations. Comparative Immunology, Microbiology and Infectious Diseases, 27(5), 357–375. doi: 10.1016/j.cimid.2004.03.005

Fyodorova, M. V., Savage, H. M., Lopatina, J. V., Bulgakova, T. A., Ivanitsky, A. V., Platonova, O. V., & Platonov, A. E. (2006). Evaluation of potential West Nile virus vectors in Volgograd Region, Russia, 2003 (Diptera: Culicidae): Species composition, bloodmeal host utilization, and virus infection rates of mosquitoes. Journal of Medical Entomology, 43(3), 552–563. doi: 10.1603/0022-2585(2006)43[552:EOPWN-V]2.0.CO;2

Gideon, B. (2014). BEDASSLE: Quantifies effects of geo/eco distance on genetic differen-tiation. Retrieved from https://cran.r-proj-ect.org/package=BEDASSLE

Gomes, B., Wilding, C. S., Weetman, D., Sousa, C. A., Novo, M. T., Savage, H. M.,… Donnelly, M. J. (2015). Limited genomic divergence between intraspecific forms of *Culex pipiens* under different ecological pressures. BMC Evolutionary Biology, 15, 197. doi: 10.1186/s12862-015-0477-z

Gordeev, M. I., & Burlak, V. A. (1991). Inversion polymorphism in malaria mosquito *Anopheles messeae*. Part X. Resistance of larvae with different genotypes to toxins of crystal-forming bacteria *Bacillus thuringiensis subsp. israelensis* (serovar H14). Genetika, 27(2), 238–246. Retrieved from http://www.ncbi.nlm.nih.gov/pubmed/1874433

Gordeev, M. I., & Moskaev, A. V. (2016). Chromosomal polymorphism in the populations of malaria mosquito *Anopheles messeae* (Diptera, Culicidae) in the Volga region. Genetika, 52(6), 685–690. Retrieved from http://www.ncbi.nlm.nih.gov/pubmed/29368498

Gordeev, M. I., & Perevozkin, V. P. (1995). Strategies for selection and stability to asphyxia in larvae of the malaria mosquito *Anopheles messeae* with various karyotypes. Genetika, 31(2), 180–184. Retrieved from http://www.ncbi.nlm.nih.gov/pubmed/7721058

Gordeev, M. I., Perevozkin, V. P., & Luk’yantsev, S. V. (1997). Genetic and ecological effects of *Argyroneta aquatica* hunting on *Anopheles* and *Culex* mosquito larvae. Genetica, 33(5), 704–709.

Gordeev, M. I., & Sibataev, A. K. (1995). Influence of the carnivorous plant pemphigus *Utricularia vulgaris* on selection processes in larvae of malarial mosquitoes. Ecology, 3, 241–245.

Gordeev, M. I., & Stegniy, V. N. (1987). Inversion polymorphism in the malaria mosquito *Anopheles messeae*. VII. Fertility and the population genetics structure of the species. Genetika, 23(12), 2169–2174. Retrieved from http://www.ncbi.nlm.nih.gov/pubmed/3440514

Gordeev, M. I., & Troshkov, N. I. (1990). Inversion polymorphism of the malaria mosquito *Anopheles messeae*. IX. Cannibalism in larvae as a selection factor. Genetika, 26(9), 1597–1603. Retrieved from http://www.ncbinlm.nih.gov/pubmed/2079206

Gornostaeva, R. M., & Danilov, A.. (2002). On distribution of malaria mosquitoes (Diptera, Culicidae: *Anopheles*) from Maculipennis Complex in Russian territory. Parasitology, 26, 33–47.

Graffelman, J. (2015). Exploring Diallelic Genetic Markers: The Hardy-Weinberg Package. Journal of Statistical Software, 64(3), 1–23. doi: 10.18637/jss.v064.i03

Gray, E. M., Rocca, K. A., Costantini, C., & Besansky, N. J. (2009). Inversion 2La is associated with enhanced desiccation resistance in *Anopheles gambiae*. Malaria Journal 8(215). doi: 10.1186/1475-2875-8-215

Gutsevich, A. V., Monchadsky, A. S., & Shtalelberg, A. A. (1970). Fauna of the U.S.S.R. Diptera (Vol. 3). Moscow: Nauka.

Haba, Y., & McBride, L. (2022). Origin and status of *Culex pipiens* mosquito ecotypes. Current Biology: CB, 32(5), R237–R246. doi: 10.1016/j.cub.2022.01.062

Hackett, L. W. (1937). Malaria in Europe: an ecological study. In London: Oxford university press.

Hackett, L. W., & Missiroli, A. (1935). The varieties of *Anopheles maculipennis* and their relation to the distribution of malaria in Europe. Riv. Malariol, 14, 45–109.

Hewitt, G. M. (2001). Speciation, hybrid zones and phylogeography - or seeing genes in space and time. Molecular Ecology, 10(3), 537–549. doi: 10.1046/J.1365-294X.2001.01202.X

Hewitt, G. M. (2004). Biodiversity: a climate for colonization. Heredity, 92(1), 1–2. doi: 10.1038/SJ.HDY.6800365

Hoffmann, A. A., Sgrò, C. M., & Weeks, A. R. (2004). Chromosomal inversion polymorphisms and adaptation. Trends in Ecology & Evolution, 19(9), 482–488. doi: 10.1016/j.tree.2004.06.013

Jones, M. D. R. (1974). Inversion polymorphism and circadian flight activity in the mosquito *Anopheles stephensi* List. (Diptera, Culicidae). Bulletin of Entomological Research, 64(2), 305–311. doi: 10.1017/S0007485300031199

Kabanova, V. M., Kartashova, N. N., & Stegniy, V. N. (1972). Karyological study of natural populations of malarial mosquitoes in the Middle Ob river. I. Characteristics of the karyotype of *Anopheles maculipennis messeae*. Tsitologiia, 14(5), 630–636. Retrieved from http://www.ncbi.nlm.nih.gov/pubmed/5035512

Kabanova, V. M., Stegniy, V. N., & Luzhkova, A. G. (1973). Seasonal dynamics of inversion polymorphism in a natural population of the malarial mosquito *Anopheles messeae*(Diptera: Culicidae). Genetika, 9(10), 78–82. Retrieved from http://www.ncbi.nlm.nih.gov/pubmed/4807110

Kamali, M., Xia, A., Tu, Z., & Sharakhov, I. V. (2012). A new chromosomal phylogeny supports the repeated origin of vectorial capacity in malaria mosquitoes of the *Anopheles gambiae* complex. PLoS Pathogens, 8(10), e1002960. doi: 10.1371/journal.ppat.1002960

Kampen, H., Schäfer, M., Zielke, D. E., & Walther, D. (2016). The *Anopheles maculipennis* complex (Diptera: Culicidae) in Germany: an update following recent monitoring activities. Parasitology Research, 115(9), 3281–3294. doi: 10.1007/s00436-016-5189-9

Kampen, H. (2005). Integration of *Anopheles beklemishevi* (Diptera: Culicidae) in a PCR assay diagnostic for palaearctic *Anopheles maculipennis* sibling species. Parasitology Research, 97(2), 113–117. doi: 10.1007/s00436-005-1392-9

Kamvar, Z. N., Tabima, J. F., & Grünwald, N. J. (2014). Poppr: an R package for genetic analysis of populations with clonal, partially clonal, and/or sexual reproduction. PeerJ, 2, e281. doi: 10.7717/peerj.281

Kapun, M., Fabian, D. K., Goudet, J., & Flatt, T. (2016). Genomic evidence for adaptive inversion clines in *Drosophila melanogaster*. Molecular Biology and Evolution, 33(5), 1317–1336. doi: 10.1093/molbev/msw016

Kavran, M., Zgomba, M., Weitzel, T., Petric, D., Manz, C., & Becker, N. (2018). Distribution of *Anopheles daciae* and other *Anopheles maculipennis* complex species in Serbia. Parasitology Research, 117(10), 3277–3287. doi: 10.1007/s00436-018-6028-y

Kitzmiller, J. B., Frizzi, G., & Baker, R. (1967). Evolution and speciation within the Maculi-pennis Complex of the genus *Anopheles*. In J. W. Wright (Ed.), Genetics of insect vectors of disease (pp. 151–210). Amsterdam-Lon-don-New York: Elsevier Publishing Company.

Kolaczkowski, B., Kern, A. D., Holloway, A. K., & Begun, D. J. (2011). Genomic differentiation between temperate and tropical Australian populations of *Drosophila melanogaster*. Genetics, 187(1), 245–260. doi: 10.1534/ge-netics.110.123059

Krimbas, C. B., & Powell, J. R. (1992). Drosophila inversion polymorphism. CRC Press.

Kronefeld, M., Werner, D., & Kampen, H. (2014). PCR identification and distribution of *Anopheles daciae* (Diptera, Culicidae) in Germany. Parasitology Research, 113(6), 2079–2086. doi: 10.1007/s00436-014-3857-1

Kulkarni, M. A., Duguay, C., & Ost, K. (2022). Charting the evidence for climate change impacts on the global spread of malaria and dengue and adaptive responses: a scoping review of reviews. Globalization and Health, 18(1), 1. doi: 10.1186/s12992-021-00793-2

Lanzaro, G. C., Touré, Y. T., Carnahan, J., Zheng, L., Dolo, G., Traoré, S.,… Taylor, C. E. (1998). Complexities in the genetic structure of *Anopheles gambiae* populations in west Africa as revealed by microsatellite DNA analysis. Proceedings of the National Academy of Sciences of the United States of America, 95(24), 14260–14265. doi: 10.1073/PNAS.95.24.14260

Lilja, T., Eklöf, D., Jaenson, T. G. T., Lindström, A., & Terenius, O. (2020). Single nucleotide polymorphism analysis of the ITS2 region of two sympatric malaria mosquito species in Sweden: *Anopheles daciae* and *Anopheles messeae*. Medical and Veterinary Entomology, 34(3), 364–368. doi: 10.1111/mve.12436

Main, B. J., Lee, Y., Ferguson, H. M., Kreppel, K.S., Kihonda, A., Govella, N. J.,… Lanzaro, G. C. (2016). The genetic basis of host preference and resting behavior in the major African malaria vector, *Anopheles arabien-sis*. PLoS Genetics, 12(9), 1–17. doi: 10.1371/journal.pgen.1006303

Mayr, E. (1963). Animal species and evolution. London: Harvard University Press.

Menhinick, E. F. (1964). A comparison of some species-individuals diversity indices applied to samples of field insects. Ecology, 45(4), 859–861. doi: 10.2307/1934933

Michalakis, Y., & Excoffier, L. (1996). A generic estimation of population subdivision using distances between alleles with special reference for microsatellite loci. Genetics, 142(3), 1061–1064. Retrieved from http://www.ncbi.nlm.nih.gov/pubmed/8849912

Naumenko, A. N., Karagodin, D. A., Yurchenko, A. A., Moskaev, A. V., Martin, O. I., Baricheva, E. M.,… Sharakhova, M. V. (2020). Chromosome and genome divergence between the cryptic Eurasian malaria vector-species *Anopheles messeae* and *Anopheles daciae*. Genes, 11(2). doi: 10.3390/genes11020165

Nicolescu, G., Linton, Y. M., Vladimirescu, A., Howard, T. M., & Harbach, R. E. (2004). Mosquitoes of the *Anopheles maculipennis* group (Diptera: Culicidae) in Romania, with the discovery and formal recognition of a new species based on molecular and morphological evidence. Bulletin of Entomological Research, 94(6), 525–535. doi: 10.1079/ber2004330

Novikov, Y. M., & Kabanova, V. M. (1979). Adaptive association of inversions in a natural population of the malaria mosquito *Anopheles messeae* Fall. Genetika, 15(6), 1033–1045. Retrieved from http://www.ncbi.nlm.nih.gov/pubmed/467975

Novikov, Y. M., & Shevchenko, A. I. (2001). Inversion polymorphism and the divergence of two cryptic forms of *Anopheles messeae* (Diptera, Culicidae) at the level of genomic DNA repeats. Russian Journal of Genetics, 37(7), 754–763. doi: 10.1023/A:1016790724790

Nwakanma, D. C., Neafsey, D. E., Jawara, M., Adiamoh, M., Lund, E., Rodrigues, A.,… Conway, D. J. (2013). Breakdown in the process of incipient speciation in *Anopheles gambiae*. Genetics, 193(4), 1221–1231. doi: 10.1534/genetics.112.148718

Oliveira, E., Salgueiro, P., Palsson, K., Vicente, J. L., Arez, A. P., Jaenson, T. G.,… Pinto, J. (2008). High levels of hybridization between molecular forms of *Anopheles gambiae* from Guinea Bissau. Journal of Medical Entomology, 45(6), 1057–1063. doi: 10.1603/0022-2585(2008)45[1057:hlohbm]2.0.co;2

Perevozkin, V. P., Printseva, A. A., Maslennikov, P. V, & Bondarchuk, S. S. (2012). Genetic aspects of sexual behavior in malaria mosquitoes on the basis of specific acoustic signals at mating. Genetika, 48(6), 692–697. Retrieved from https://pubmed.ncbi.nlm.nih.gov/22946326/

Petrarca, V., & Beier, J. C. (1992). Intraspecific chromosomal polymorphism in the *Anopheles gambiae* complex as a factor affecting malaria transmission in the Kisumu area of Kenya. The American Journal of Tropical Medicine and Hygiene, 46(2), 229–237. doi: 10.4269/ajtmh.1992.46.229

Pleshkova, G. N., Stegniy, V. N., Novikov, Y. M., & Kabanova, V. M. (1978). Inversion polymorphism in the malarial mosquito, *Anopheles messeae*. III. The temporal dynamics of the inversion frequencies in the population of the center of a geographic range. Genetika, 14, 2169–2176.

Powell, J. R. (2018). Genetic Variation in Insect Vectors: Death of Typology? Insects, 9(4). doi: 10.3390/insects9040139

Proft, J., Maier, W. A., & Kampen, H. (1999). Identification of six sibling species of the *Anopheles maculipennis* complex (Diptera: Culicidae) by a polymerase chain reaction assay. Parasitology Research, 85(10), 837–843. doi: 10.1007/s004360050642

R_Core_Team. (2021). R: A language and environment for statistical computing. Vienna: R Foundation for Statistical Computing. Retrieved from https://www.r-project.org/

Rieseberg, L. H. (2001). Chromosomal rearrangements and speciation. Trends in Ecology & Evolution, 16(7), 351–358. doi: 10.1016/s0169-5347(01)02187-5

Rossati, A., Bargiacchi, O., Kroumova, V., Zaramella, M., Caputo, A., & Garavelli, P. L. (2016). Climate, environment and transmission of malaria. Le Infezioni in Medicina, 24(2), 93–104. doi: 10.1371/currents.outbreaks.95fbc4a8fb4695e049baabfc2fc8289f

Rougeron, V., Boundenga, L., Arnathau, C., Durand, P., Renaud, F., & Prugnolle, F. (2022). A population genetic perspective on the origin, spread and adaptation of the human malaria agents *Plasmodium falciparum* and *Plasmodium vivax*. FEMS Microbiology Reviews, 46(1). doi: 10.1093/FEMSRE/FUAB047

Rydzanicz, K., Czułowska, A., Manz, C., & Jawień, P. (2017). First record of *Anopheles daciae* (Linton, Nicolescu & Harbach, 2004) in Poland. Journal of Vector Ecology: Journal of the Society for Vector Ecology, 42(1), 196–199. doi: 10.1111/jvec.12257

Sainz-Elipe, S., Latorre, J. M., Escosa, R., Masià, M., Fuentes, M. V., Mas-Coma, S., & Bargues, M. D. (2010). Malaria resurgence risk in southern Europe: climate assessment in an historically endemic area of rice fields at the Mediterranean shore of Spain. Malaria Journal, 9, 221. doi: 10.1186/1475-2875-9-221

Schaffner, F., Marquine, M., Pasteur, N., & Raymond, M. (2003). Genetic differentiation of *Anopheles claviger s.s*. in Europe. Journal of Medical Entomology, 40(6), 865–875. doi: 10.1603/0022-2585-40.6.865

Shannon, C. E. (1948). A mathematical theory of communication. Bell System Technical Journal, 27(3), 379–423. doi: 10.1002/j.1538-7305.1948.tb01338.x

Sharakhov, I. V., White, B. J., Sharakhova, M. V., Kayondo, J., Lobo, N. F., Santolamazza, F.,… Besansky, N. J. (2006). Breakpoint structure reveals the unique origin of an interspecific chromosomal inversion (2La) in the *Anopheles gambiae* complex. Proceedings of the National Academy of Sciences of the United States of America, 103(16), 6258–6262. doi: 10.1073/pnas.0509683103

Simard, F., Ayala, D., Kamdem, G. C., Pombi, M., Etouna, J., Ose, K.,… Costantini, C. (2009). Ecological niche partitioning between *Anopheles gambiae* molecular forms in Cameroon: the ecological side of speciation. BMC Ecology, 9. doi: 10.1186/14726785-9-17

Simpson, E. H. (1949). Measurement of Diversity. Nature, 163(4148), 688–688. doi: 10.1038/163688a0

Sinka, M. E., Bangs, M. J., Manguin, S., Coetzee, M., Mbogo, C. M., Hemingway, J.,… Hay, S. I. (2010). The dominant *Anopheles* vectors of human malaria in Africa, Europe and the Middle East: occurrence data, distribution maps and bionomic précis. Parasites & Vectors, 3, 117. doi: 10.1186/1756-3305-3-117

Smitz, N., De Wolf, K., Gheysen, A., Deblauwe, I.,Vanslembrouck, A., Meganck, K.,… Van Bortel, W. (2021). DNA identification of species of the *Anopheles maculipennis* complex and first record of *An. daciae* in Belgium. Medical and Veterinary Entomology, 35(3), 442–450. doi: 10.1111/mve.12519

Stegniy, V. N. (1976). Revealing of chromosme races in the malaria mosquito *Anopheles sacharovi* (Diptera, Culicidae). Tsitologia, 18. 1039–1041

Stegniy, V. N. (1980). Reproductive interrelations of malarial mosquitoes of the complex *Anopheles maculipennis* (Diptera, Culicidae) Zoologicheskii Zhurnal, 59(10) 1469–1475

Stegniy, V. N. (1983a). Inversion polymorphism of the malarial mosquito *Anopheles messeae*. IV. The stability of the frequency distribution of the inversions by species area. Genetika, 19, 466–473.

Stegniy, V. N. (1983b). Inversion polymorphism of the malarial mosquito *Anopheles messeae*. V. The interaction of different chromosomal inversions in the spatial area. Genetica, 19, 474–482.

Stegniy, V. N. (1991). Population Genetics and Evolution of Malaria Mosquitoes. Tomsk: Tomsk State University Publisher.

Stegniy, V. N., & Kabanova, V. M. (1978). Cy-toecological study of natural populations of malaria mosquitoes on the USSR territory. 1. Isolation of a new species of *Anopheles* in Maculipennis complex by the cytodiagnos-tic method. Meditsinskaia Parazitologiia i Parazitarnye Bolezni, 45(2), 192–198. Retrieved from http://www.ncbi.nlm.nih.gov/pubmed/1024140

Stegniy, V. N., Kabanova, V. M., & Novikov, Y. M. (1976). Study of the karyotype of the malaria mosquito. Tsitologiia, 18(6), 760–766.

Stegniy, V. N., Kabanova, V. M., Novikov, Y. M., & Pleshkova, G. N. (1976). Inversion polymorphism in malaria mosquito *Anopheles messeae*. I. Distribution of the inversions in the species areal. Genetika, 12, 47–55.

Stegniy, V. N., Novikov, Y. M., Pleshkova, G. N., & Kabanova, V. M. (1978). Inversion polymorphism in malaria mosquito *Anopheles messeae*. II. Interpopulation variablity of inversion frequncies. Russian Journal of Genetics, 14, 1017–1023.

Stegniy, V. N., Pestryakova, T. S., & Kabanova, V. M. (1973). Cytological identification of sibling species of the malaria mosquitoes *Anopheles maculipennis* and *An. messeae*. Zool. Zhurnal, 52, 1971–1976.

Stegniy, V. N., Pishchelko, A. O., Sibataev, A. K., & Abylkassymova, G. (2016). Spatial and temporal variations of the chromosomal inversion frequencies across the range of malaria mosquito *Anopheles messeae* Fall. (Culicidae) during the 40-year monitoring period. Genetika, 52(6), 664–671. Re-trieved from http://www.ncbi.nlm.nih.gov/pubmed/29368494

Stegniy, V. N., Sichinava, S. G., & Sipovich, N. G. (1984). A hybridological analysis and biology of the malarial mosquitos of the complex *Anopheles maculipennis* (Diptera, Culicidae) in Western Georgia. Zoologichesky Zhurnal, 63, 300–303.

Torsvik, T. H., & Cocks, R. M. (2017). Earth history and palaeogeogrphy. Cambridge, UK: Cambridge University Press.

Touré, Y. T. (1989). The current state of studies of malaria vectors and the antivectorial campaign in west Africa. Transactions of the Royal Society of Tropical Medicine and Hygiene, 83 Suppl, 39–41. doi: 10.1016/0035-9203(89)90602-0

Touré, Y. T., Petrarca, V., Traoré, S. F., Coulibaly, A., Maïga, H. M., Sankaré, O.,… Coluzzi, M. (1994). Ecological genetic studies in the chromosomal form Mopti of *Anopheles gambiae* s.str. in Mali, west Africa. Genetica, 94(2–3), 213–223. doi: 10.1007/BF01443435

Tzedakis, P. C., Emerson, B. C., & Hewitt, G. M. (2013). Cryptic or mystic? Glacial tree refugia in Northern Europe. Trends in Ecology & Evolution, 28(12), 696–704. doi: 10.1016/J.TREE.2013.09.001

Vaulin, O. V., & Novikov, Y. M. (2012). Polymorphism and interspecific variability of cytochrome oxydase subunit I (COI) gene nucleotide sequence in sibling species of A and B Anopheles messeae and *An. beklemishevi* (Diptera: Culicidae). Vavilov Journal of Genetics and Breeding, 16(2), 358–368.

Venkatesan, M., Westbrook, C. J., Hauer, M. C., & Rasgon, J. L. (2007). Evidence for a population expansion in the West Nile virus vector *Culex tarsalis*. Molecular Biology and Evolution, 24(5), 1208–1218. doi: 10.1093/MOLBEV/MSM040

Weir, B. S., & Hill, W. G. (2002). Estimating F-statistics. Annual Review of Genetics, 36, 721–750. doi: 10.1146/annurev.gen-et.36.050802.093940

Weitzel, T., Gauch, C., & Becker, N. (2012). Identification of *Anopheles daciae* in Germany through ITS2 sequencing. Parasitology Research, 111(6), 2431–2438. doi: 10.1007/s00436-012-3102-8

Wigginton, J. E., Cutler, D. J., & Abecasis, G. R. (2005). A note on exact tests of Hardy-Weinberg equilibrium. American Journal of Human Genetics, 76(5), 887–893. doi: 10.1086/429864

Winter, D. J. (2012). MMOD: an R library for the calculation of population differentiation statistics. Molecular Ecology Resources, 12(6), 1158–1160. doi: 10.1111/j.1755-0998.2012.03174.x

Yurchenko, A. A., Masri, R. A., Khrabrova, N. V, Sibataev, A. K., Fritz, M. L., & Sharakhova, M. V. (2020). Genomic differentiation and intercontinental population structure of mosquito vectors *Culex pipiens pipiens* and *Culex pipiens molestus*. Scientific Reports, 10(1), 7504. doi: 10.1038/s41598-020-63305-z

Yurchenko, A. A., Naumenko, A. N., Artemov G. N., Karagodin, D. A., Hodge, J. M., Velichevskaya, A. I., Kokhanenko, A. A., Bondarchuk, S. S., Abai, M. R., Kamali, M., Gordeev, M. I., Moskaev, A. V.,. Caputo, B, Aghayan, S. A., Baricheva, E. M., Stegniy, V. N., Sharakhova, M. V., and Sharakhov, I. V. (2022). Phylogenomics revealed migration routes and adaptive radiation timing of Holarctic malaria vectors of the Maculipennis group. bioRxiv 2022.08.10.503503.

Zvantsov, A. B., Gordeev, M. I., Goriacheva, I. I., & Ezhov, M. N. (2014). The distribution of the mosquitoes of the *Anopheles maculipennis* complex (Diptera, Culicidae, Anophelinae) in Central Asia. Meditsinskaia Parazitologiia i Parazitarnye Bolezni, (4), 19–23. Retrieved from http://www.ncbi.nlm.nih.gov/pubmed/25812402

